# The timing and impact of psychiatric, cognitive and motor abnormalities in Huntington’s disease

**DOI:** 10.1101/2020.05.26.116798

**Authors:** Branduff McAllister, James F. Gusella, G. Bernhard Landwehrmeyer, Jong-Min Lee, Marcy E. MacDonald, Michael Orth, Anne E. Rosser, Nigel M. Williams, Peter Holmans, Lesley Jones, Thomas H. Massey, on behalf of the REGISTRY Investigators of the European Huntington’s disease network

## Abstract

**Objective:** To assess the prevalence, timing and functional impact of psychiatric, cognitive and motor abnormalities in Huntington’s disease (HD) gene carriers, we analysed retrospective clinical data from individuals with manifest HD.

**Methods:** Clinical features of HD patients were analysed for 6316 individuals in the European REGISTRY study from 161 sites across 17 countries. Data came from clinical history and the patient-completed Clinical Characteristics Questionnaire that assessed eight symptoms: motor, cognitive, apathy, depression, perseverative/obsessive behavior, irritability, violent/aggressive behavior, and psychosis. Multiple logistic regression was used to analyse relationships between symptoms and functional outcomes.

**Results:** The initial manifestation of HD is increasingly likely to be motor, and less likely to be psychiatric, as age at presentation increases, and is independent of pathogenic CAG repeat length. The Clinical Characteristics Questionnaire captures data on non-motor symptom prevalence that correlate specifically with validated clinical measures. Psychiatric and cognitive symptoms are common in HD gene carriers, with earlier onsets associated with longer CAG repeats. 42.4% of HD patients reported at least one psychiatric or cognitive symptom before motor symptoms, with depression most common. Each non-motor symptom was associated with significantly reduced total functional capacity scores.

**Conclusions:** Psychiatric and cognitive symptoms are common and functionally debilitating in HD gene carriers. They require recognition and targeting with clinical outcome measures and treatments. However, as it is impossible to distinguish confidently between non-motor symptoms arising from HD and primary psychiatric disorders, particularly in younger pre-manifest patients, non-motor symptoms should not be used to make a clinical diagnosis of HD.

## Introduction

Huntington’s disease (HD) is a central neurodegenerative disorder caused by an expanded CAG repeat (>35 CAGs) in the *Huntingtin* gene^1^. Longer repeats are associated with earlier disease onset^2, 3^. Neuronal loss in the brain causes progressive motor abnormalities, cognitive decline and ultimately death. The movement disorder usually includes chorea, but may also involve dystonia, ataxia, oculomotor problems and parkinsonism, some of which are initially only identifiable through targeted HD examination. Debilitating behavioral and psychiatric symptoms are common in HD gene carriers, and require treatment, although they cannot be used in clinical practice to define HD onset because it is impossible to distinguish psychiatric manifestations of HD from coincident diagnoses^4, 5^. Prospective studies of HD gene carriers many years from predicted clinical onset have shown only subtle motor, cognitive and psychiatric deficits when compared with age and sex-matched controls^6–8^. This implies there is a window for therapeutic intervention to preserve normal brain functions. Understanding in detail the timing and impact of different symptoms in HD gene carriers will help improve targeted therapies.

The HD Clinical Characteristics Questionnaire (HD-CCQ)^9^ gathers retrospective data from individuals with HD about the prevalence and timing of eight motor, cognitive and psychiatric symptoms^10^. Here we validate HD-CCQ data for non-motor symptoms by showing strong and specific associations with established scores of depression, irritability and cognition. We use HD-CCQ data to show the high prevalence of psychiatric and cognitive symptoms in HD gene carriers, often in advance of motor symptoms, and their negative impact on the lives of patients.

## Methods

### Standard protocol approvals, registrations and patient consents

Participants were in the multicentre, multinational, observational REGISTRY study of European HD (http://www.ehdn.org/wp-content/uploads/2018/06/registry-protocol-3.0.pdf; <u>NCT01590589</u>). Data were accessed as part of European Huntington’s Disease Network (EHDN) data mining project 0791. Ethical approval for REGISTRY was obtained in each participating country. All participants gave written informed consent.

### Participant data

HD participant data, collected June 2004 - February 2016 across 161 sites in 17 European countries, was obtained for 6316 individuals (accessed October 2016) who had clinical HD onset, determined by the rating clinician in REGISTRY, and a confirmed pathogenic CAG length of 36-93 CAGs. Of these CAG sizes, 5027 were centrally determined by BioRep Inc. (Milan, Italy; REGISTRY protocols) and 1289 were derived by local diagnostic laboratories. Two estimates of the age at onset of symptoms and/or signs in HD were used in this study. First, the clinician-estimated age at first HD manifestation based upon all available clinical evidence at the first REGISTRY visit (coded as ‘sxrater’). Having a ‘sxrater’ age at onset was required for inclusion in this study. Onset type was classified as motor, cognitive, psychiatric, oculomotor, other or mixed. As the clinician’s estimate was given as a date, age estimates were calculated using the participant’s anonymised birthday; where only a year was given, July 15^th^ was used for estimation (15/07/xxxx). Second, the ages at onset of different symptoms in HD patients were estimated by the HD Clinical Characteristics Questionnaire (HD-CCQ) which was completed by a healthcare professional, usually a HD-specialist nurse or similarly-qualified person, using responses from the individual with HD and their care partners (present in clinic in 93.1% of cases), and patient medical notes. The HD-CCQ comprises questions about eight symptoms commonly observed in HD, asking whether the participant has ever had the symptom (yes or no), and, if yes, the age at which the symptom was first experienced (Appendix 3). Information was available, at least in part, for 5609 individuals. The symptoms recorded (number of individuals with data) were: motor (chorea or other, consistent with HD; 5603), cognitive impairment sufficient to impact on work or daily living (5591), apathy (5584), depression (5595), perseverative/obsessive behaviour (5588), irritability (5586), violent or aggressive behaviour (5586) and psychosis (5589). For subsequent analyses, missing data were handled using pairwise deletion to maximise the number of individuals. Typically, the rater estimate of clinical onset and initial HD-CCQ would be recorded at the first REGISTRY visit, sometimes by one clinician, and sometimes by a clinician and another qualified staff member such as HD-specialist nurse, depending on local clinic set-up. Subsequent visits updated the HD-CCQ: we used data from the most recent clinic visit. We had data on Shoulson-Fahn disease stage at last clinic visit for 4554 individuals (72.1% of our study population): stage 1 (Total Functional Capacity (TFC) 11-13; N=890; 19.5%), stage 2 (TFC 7-10; N=1278; 28.1%), stage 3 (TFC 4-6; N=969; 21.3%), stage 4 (TFC 1-3; N=1133; 24.9%), stage 5 (TFC 0; N=284; 6.2%).

The Hospital Anxiety/Depression Scale (HADS) and Snaith Irritability Scale (SIS) were completed by the participant at each clinic visit and provide measures of anxiety, depression and irritability at that specific time. We used lifetime highest total depression and total irritability scores from both the HADS and the SIS in analyses. Similarly, the symbol-digit modalities test (SDMT) and Stroop tests of cognitive ability were administered as part of the Unified Huntington’s Disease Rating Scale (UHDRS)^11^ at each visit. The UHDRS consists of validated questionnaires, tools and examinations related to motor, cognitive, behavioural and functional impairments seen in HD. For the SDMT and Stroop, we used the total correct scores from the most recent clinic visit. Disease duration was estimated by taking the most recent visit and subtracting the clinician’s estimate of disease onset. The product of Problem Behaviours Assessment (PBA-s) severity and frequency scores from the most recent clinic was used for modelling purposes.

### Statistical analyses of clinical data

Total depression scores from HADS, total irritability scores from SIS, the number of correct answers in the SDMT, the number of correct answers in Stroop tests or composite PBA-s scores were regressed on HD clinical characteristics data, age, CAG length, sex and disease duration (table 1). To calculate coefficients of determination (R^2^ values, table 2), HD-CCQ age at onset data were natural log transformed. Only individuals with a known sex and a symptom onset ≥3 years were considered, and a residual vs leverage plot identified one influential data point passing Cook’s distance that was removed from all R^2^ calculations. *P* values were calculated comparing male and female R^2^ values using Fisher’s transformation^12^. A chi-square test was used to test for differences in symptom frequency, derived from the yes/no component of the HD-CCQ, between males and females.

**Table 1.** Association of validated clinical scores with the HD Clinical Characteristics Questionnaire symptoms (shown in italics), and other covariates. For binary covariates (CCQ symptoms and sex) “effect” is the increase/decrease in the clinical score associated with presence of that covariate. For quantitative covariates (age, CAG, duration), “effect” is the change in clinical score associated with an increase of one unit in the covariate. In addition to having a confirmed onset and pathogenic CAG length (36-93), individuals must have no co-morbid diagnosis of schizophrenia, schizotypy or schizoaffective disorder. Significant associations after Bonferroni correction for 4 phenotypes and 12 covariates are shown in bold (*P* < 1.04 x 10^-3^) and nominally significant P values are italicised (*P* < 0.05). CI: confidence interval; TDS: Total depression score from the Hospital Anxiety and Depression Scale; TIS: Total irritability score from Snaith’s irritability scale; SDMT: Symbol digits modalities test; POB: perseverative/obsessive behaviour; VAB: violent or aggressive behaviour; Sex (F): female.

|  | TDS (N=2403) |  | TIS (N=2403) |  | SDMT (N=3137) |  | Stroop Interference (N=3273) |  |
| --- | --- | --- | --- | --- | --- | --- | --- | --- |
|  | Effect (95% CI) | *P* | Effect (95% CI) | *P* | Effect (95% CI) | *P* | Effect (95% CI) | *P* |
| *Motor* | 0.15 (±1.85) | 8.71 x 10<sup>-1</sup> | 0.67 (±1.93) | 4.93 x 10<sup>-1</sup> | -12.37 (±3.90) | **5.62 x 10<sup>-10</sup>** | -9.34 (±3.76) | **1.19 x 10<sup>-6</sup>** |
| *Cognitive* | 0.38 (±0.39) | 5.28 x 10<sup>-2</sup> | -0.41 (±0.40) | 4.53 x 10<sup>-2</sup> | -3.52 (±0.81) | **2.28 x 10<sup>-17</sup>** | -3.41 (±0.76) | **3.29 x 10<sup>-18</sup>** |
| *Apathy* | 1.73 (±0.40) | **4.05 x 10<sup>-17</sup>** | 0.48 (±0.42) | 2.36 x 10<sup>-2</sup> | -2.70 (±0.83) | **2.37 x 10<sup>-10</sup>** | -2.15 (±0.79) | **1.00 x 10<sup>-7</sup>** |
| *Depression* | 1.49 (±0.40) | **5.67 x 10<sup>-13</sup>** | 1.13 (±0.42) | **1.37 x 10<sup>-7</sup>** | -0.09 (±0.84) | 8.41 x 10<sup>-1</sup> | -0.53 (±0.79) | 1.88 x 10<sup>-1</sup> |
| *POB* | -0.15 (±0.42) | 4.96 x 10<sup>-1</sup> | 0.04 (±0.44) | 8.70 x 10<sup>-1</sup> | -1.28 (±0.85) | 3.26 x 10<sup>-3</sup> | -1.10 (±0.80) | 7.45 x 10<sup>-3</sup> |
| *Irritability* | 0.15 (±0.43) | 4.97 x 10<sup>-1</sup> | 1.82 (±0.45) | **1.99 x 10<sup>-15</sup>** | 1.28 (±0.89) | 4.52 x 10<sup>-3</sup> | 1.01 (±0.84) | 1.76 x 10<sup>-2</sup> |
| *VAB* | 0.72 (±0.47) | 2.65 x 10<sup>-3</sup> | 1.57 (±0.49) | **3.29 x 10<sup>-10</sup>** | -1.24 (±0.97) | 1.26 x 10<sup>-2</sup> | -1.20 (±0.92) | 1.06 x 10<sup>-2</sup> |
| *Psychosis* | -0.45 (±0.68) | 1.98 x 10<sup>-1</sup> | -0.50 (±0.71) | 1.67 x 10<sup>-1</sup> | -2.53 (±1.38) | **3.38 x 10<sup>-4</sup>** | -3.18 (±1.28) | **1.07 x 10<sup>-6</sup>** |
| Age | 0.01 (±0.02) | 2.18 x 10<sup>-1</sup> | -0.09 (±0.02) | **8.17 x 10<sup>-13</sup>** | -0.53 (±0.05) | **1.02 x 10<sup>-99</sup>** | -0.52 (±0.05) | **2.89 x 10<sup>-104</sup>** |
| CAG | -0.03 (±0.06) | 3.54 x 10<sup>-1</sup> | -0.21 (±0.07) | **1.17 x 10<sup>-9</sup>** | -1.56 (±0.14) | **3.97 x 10<sup>-102</sup>** | -1.25 (±0.13) | **2.38 x 10<sup>-71</sup>** |
| Sex (F) | -0.12 (±0.37) | 5.17 x 10<sup>-1</sup> | 0.35 (±0.38) | 7.14 x 10<sup>-2</sup> | -1.28 (±0.76) | **9.73 x 10<sup>-4</sup>** | -1.22 (±0.72) | **9.13 x 10<sup>-4</sup>** |
| Duration | 0.05 (±0.04) | 5.62 x 10<sup>-3</sup> | 0.02 (±0.04) | 3.08 x 10<sup>-1</sup> | -0.41 (±0.08) | **4.78 x 10<sup>-25</sup>** | -0.32 (±0.07) | **4.82 x 10<sup>-18</sup>** |

**Table 2.** Lifetime prevalence of motor and psychiatric symptoms in males and females with HD. Data from HD Clinical Characteristics Questionnaire at last recorded clinic visit in Registry. Chi-square (c^2^) tests the difference between prevalence in males and females. Odds ratios (OR) >1 indicates the symptom is more common in females; odds ratios <1 indicates the symptom is more common in males. To be included, individuals must have a pathogenic CAG length (36-93) and confirmed clinical HD onset. Significant *P* values in bold (*P* < 6.25 x 10^-3^, multiple testing correction). CI: confidence interval; POB: perseverative/obsessive behaviour; VAB: violent or aggressive behaviour.

|  | Males |  |  | Females |  |  | OR<br>(95% CI) | P value (c<sup>2</sup>) |
| --- | --- | --- | --- | --- | --- | --- | --- | --- |
|  | Yes | No | Frequency | Yes | No | Frequency |  |  |
| Motor | 2691 | 28 | 98.97% | 2859 | 25 | 99.13% | 1.19 (0.69 - 2.05) | 5.29 x 10<sup>-1</sup> |
| Cognitive | 1584 | 1132 | 58.32% | 1688 | 1187 | 58.71% | 1.02 (0.91 - 1.13) | 7.66 x 10<sup>-1</sup> |
| Apathy | 1456 | 1259 | 53.63% | 1495 | 1374 | 52.11% | 0.94 (0.85 - 1.05) | 2.56 x 10<sup>-1</sup> |
| Depression | 1582 | 1135 | 58.23% | 2025 | 853 | 70.36% | 1.70 (1.52 - 1.90) | **2.57 x 10<sup>-21</sup>** |
| POB | 1005 | 1711 | 37.00% | 1038 | 1834 | 36.14% | 0.96 (0.86 - 1.07) | 5.04 x 10<sup>-1</sup> |
| Irritability | 1706 | 1006 | 62.91% | 1634 | 1240 | 56.85% | 0.78 (0.70 - 0.87) | **4.03 x 10<sup>-6</sup>** |
| VAB | 947 | 1769 | 34.87% | 777 | 2100 | 27.01% | 0.69 (0.62 - 0.77) | **1.99 x 10<sup>-10</sup>** |
| Psychosis | 319 | 2396 | 11.75% | 325 | 2549 | 11.31% | 0.96 (0.81 - 1.13) | 6.06 x 10<sup>-1</sup> |

Associations between binary responses in the HD-CCQ (1; experienced the symptom and 0; symptom not experienced) and clinical covariates were tested using logistic regression. The covariates used were sex, CAG length, alcohol consumption (units per week), tobacco use (cigarettes per day), education (years of education), TFC score and total motor score (TMS). An additional analysis regressed the type of HD onset defined by the clinician, coded as a binary variable, on the clinician’s onset or CAG length (table e-2, doi:10.5061/dryad.pk0p2ngkz). This analysis was restricted to HD participants with CAGs 36-59, to be consistent with figure 1 subgroups, and also to adult-onset HD individuals (≥20 years). We also tested whether symptom presence was associated with the length of the wild-type (6-35 CAGs) and expanded CAGs (36-93 CAGs) alleles in individuals of known sex, and for whom both CAG lengths were known (table e-3, doi:10.5061/dryad.pk0p2ngkz). 19 individuals with a coincident formal diagnosis of schizophrenia, schizotypal disorder or schizoaffective disorder (ICD-10 F20, F21 or F25) were excluded from all models, although it was not possible to formally exclude these symptoms being part of the HD phenotype. Statistical analysis used *R* (version 3.6.0; R core team, 2019, https://www.r-project.org/).

**Figure 1.**
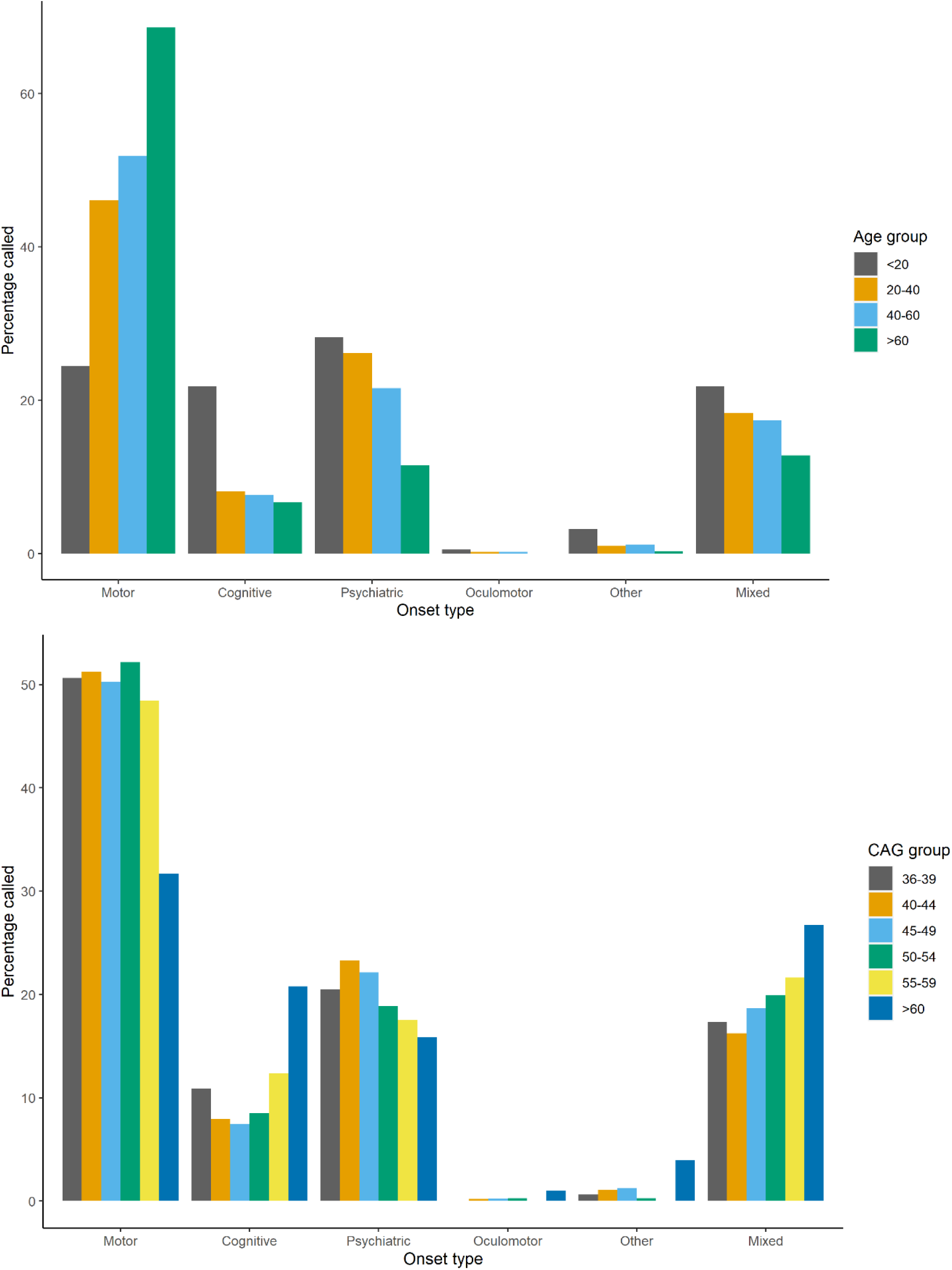
The initial manifestation of HD varies with age and CAG length. All included individuals had a pathogenic CAG length (36-93) and confirmed HD onset age determined by a rating clinician. **(A)** Frequency of different onset types in four age groups, chosen to show juvenile HD and then 20 year bins for clarity. Total N=6289; <20 years, N=188; 20-40 years, N=2216; 40-60 years, N=3276; >60 years, N=609. **(B)** Frequency of different onset types in six CAG length groups, chosen for clarity across the pathogenic range. Total N=6289; 36-39 CAG, N=156; 40-44 CAG, N=3813; 45-49 CAG, N=1735; 50-54 CAG, N=387; 55-59 CAG, N=97; >60 CAG, N=101.

### Data availability

Further information and data requests should be directed to Thomas H. Massey. Anonymised summary data is available to qualified investigators. Furthermore, anonymised patient data is available from the European Huntington’s Disease Network (EHDN) upon request given institutional assurance patient confidentiality will be upheld, and no attempt will be made to discover the identity of patients.

## Results

### The initial manifestation of HD varies with age and CAG length

The age at onset of the first unequivocal motor features of HD (‘motor onset’) has been used as a specific milestone in the natural history of HD in individuals, although it is only a crude measure of a progressive neuropathological process. It has proven particularly useful in recent genetic modifier studies of HD^13, 14^. The first psychiatric and cognitive manifestations of HD are more difficult to define with certainty, being less specific for HD and clinically indistinguishable from common coincident psychiatric diagnoses (e.g. depression), particularly in younger patients many years from predicted motor onset. The timing of the first unequivocal feature of HD is typically retrospectively recorded by a rating physician in observational studies such as REGISTRY, based on clinical information and symptom history from patients and care partners^9, 15, 16^. The rater also records the initial major presenting feature out of a choice of six: motor, cognitive, psychiatric, oculomotor, other or mixed. We analysed the initial manifestation of HD for 6316 participants in REGISTRY^9^, including 3083 males (48.8%) and 3233 females (51.2%). All participants had a confirmed genetic diagnosis of HD with a pathogenic CAG repeat length of 36-93 (figure e-1, doi:10.5061/dryad.pk0p2ngkz). The first manifestation of HD, determined by the rating physician, varied with patient age (figure 1A and table e-1, doi:10.5061/dryad.pk0p2ngkz). Individuals with onset before the age of 20, defined as juvenile HD, were equally likely to present with motor (24.5%), cognitive (21.8%) or psychiatric features (28.2%). In contrast, the initial manifestation of HD was more likely to be motor than psychiatric in adult-onset HD. As age at first manifestation increased (figure 1A and table e-2A, doi:10.5061/dryad.pk0p2ngkz) motor presentations became more likely (odds ratio (OR) = 1.06 per ten year increase in onset age, 95% confidence interval (CI) 1.04-1.07; *P* = 7.4 x 10^-22^) but psychiatric presentations became less likely (OR = 0.96 per ten year increase in onset age, 95% CI 0.95-0.97; *P* = 9.4 x 10^-16^). For people presenting over the age of 60, over two-thirds (68.6%) had initial motor abnormalities with far fewer having psychiatric (11.5%) or cognitive (6.7%) presentations. Next, we tested whether there was any relationship between pathogenic CAG repeat length, known to be inversely correlated with age at clinical onset, and the presenting phenotype. Interestingly, there was no significant relationship between CAG length (36-59 inclusive) and the relative proportions of motor, cognitive and psychiatric onset cases (figure 1B and table e-2B, doi:10.5061/dryad.pk0p2ngkz). For the few cases with data and repeat lengths of more than 59 CAG we observed a more balanced distribution of motor, cognitive and psychiatric presentations, mirroring the trends seen for the juvenile HD cases.

### Psychiatric and cognitive symptoms captured by the Clinical Characteristics Questionnaire correlate with scores from validated clinical tools

The HD Clinical Characteristics Questionnaire (HD-CCQ) was introduced to later versions of REGISTRY as the best retrospective way of capturing symptom data in existing HD populations. It is completed by a healthcare professional using information from individuals with HD and their care partners, present in clinic for over 93%, about lifetime history and age at onset of eight symptoms typical of HD. These symptoms are motor (compatible with HD), depression, irritability, violent or aggressive behavior, apathy, perseverative/obsessive behavior, psychosis and cognitive impairment sufficient to impact on work or daily living. In REGISTRY, this information was updated at each annual clinic visit. In HD-CCQ, motor symptoms are not specified beyond being compatible with HD, limiting the utility of motor data, but psychiatric and behavioral symptoms are clearly defined.

Since prevalence data from HD-CCQ have not been used in large analyses before we first tested how well they correlated with validated clinical scores of depression (HADS), irritability (SIS) and cognition (SDMT and Stroop). To mitigate against potential effects of medication at certain times, we used the lifetime highest total depression and total irritability scores for each individual. For cognitive tests we used scores at the last recorded clinic visit as these would be expected to worsen progressively and be little affected by medication. Total depression score from HADS was significantly increased in individuals with depression recorded in HD-CCQ (table 1; increase of 1.49 units, 95% CI 1.09-1.89; *P* = 5.7 x 10^-13^). An increase in HADS score was also observed in individuals with HD-CCQ apathy, probably because apathy, common in HD, may be mistaken for depression by individuals and their care partners when completing the HD-CCQ. Total irritability score from SIS was significantly increased in individuals with HD-CCQ irritability (increase of 1.82 units, 95% CI 1.37-2.27; *P* = 2.0 x 10^-15^), and also with violent/aggressive behaviour (increase of 1.57 units, 95% CI 1.08-2.06; *P* = 3.3 x 10^-10^), as expected. Both SDMT and Stroop scores of cognitive ability were significantly decreased in individuals with cognitive impairment as recorded in HD-CCQ (reductions of 3.52 units, 95% CI 2.71-4.33; *P* = 2.3 x 10^-17^ and 3.41 units, 95% CI 2-65-4.17; *P* = 1.4 x 10^-22^, respectively). Significant associations between cognitive scores and motor and apathy symptoms were also observed. In addition, we found robust and specific associations between neuropsychiatric symptoms recorded in HD-CCQ and their related symptoms scored using the validated short-form Problem Behaviors Assessment (PBA-s; supplemental table e-4, doi:10.5061/dryad.pk0p2ngkz). The specificity of the associations between HD-CCQ data and recognised clinical scales validated the use of HD-CCQ data in subsequent analyses.

### Psychiatric symptoms are common in HD gene carriers and are associated with CAG repeat length

We next analysed the lifetime prevalence of the eight symptoms recorded in HD-CCQ in 5609 individuals with HD at their most recent clinic visit (table 2). The mean age at last recorded clinic visit was 53.3 years; 53.5 years for males with data (range 10.4 – 92.6 years; N = 2569) and 53.2 years for females (range 7.9 – 90.2 years; N = 2698). Almost all (>99%) had experienced motor symptoms compatible with HD, indicating why motor abnormalities remain the diagnostic standard for clinical onset of HD. Although motor symptoms are not defined explicitly in HD-CCQ, contemporaneous data from UHDRS showed that 96.8% of our study population had chorea, alongside variable amounts of incoordination, dystonia and rigidity. In HD gene carriers these motor symptoms are likely to be specific manifestations of HD. The next most prevalent symptom was depression, occurring in 64.5% of HD individuals with significantly more females affected than males (70.4% vs 58.2%; OR 1.70, 95% CI 1.52-1.90; *P* = 2.6 x 10^-21^). Cognitive impairment sufficient to impact upon work or activities of daily living, apathy and irritability were also each observed in over half of our HD population. Cognitive impairment and apathy were equally likely in males and females, but there was significantly more irritability observed in males (62.9% vs 56.9%; OR 0.78, 95% CI 0.70-0.87; *P* = 4.0 x 10^-6^). An excess of violent or aggressive behaviour was also observed in the male group (34.9% vs 27.0%; OR 0.69, 95% CI 0.62-0.77; *P* = 2.0 x 10^-10^). Psychosis was the least prevalent of the eight recorded symptoms, although this was still observed in over 11% of individuals with HD with no significant difference in prevalence between males and females.

There was a strong inverse correlation between pathogenic CAG repeat length (40-55 CAG inclusive) and mean age at symptom onset for all symptoms analysed (figure 2). We found no effect of wild-type CAG allele length on any symptom onset, nor any significant statistical interaction between expanded and wild-type repeat lengths (table e-3, doi:10.5061/dryad.pk0p2ngkz). Pathogenic CAG length explained 66.3% of the variance in age at onset of motor symptoms, in line with previous estimates^2, 3, 17–23^, but also between 37.5% and 61.9% of the variance in onset of each of the psychiatric symptoms analysed (table 2). Depression had the weakest association with CAG repeat length (R^2^ = 37.5%). CAG length accounted for significantly more of the variance in age at onset of perseverative/obsessive behaviour in males (table 2; *P* = 3.7 x 10^-3^) and irritability in females (*P* = 1.3 x 10^-3^).

**Figure 2.**
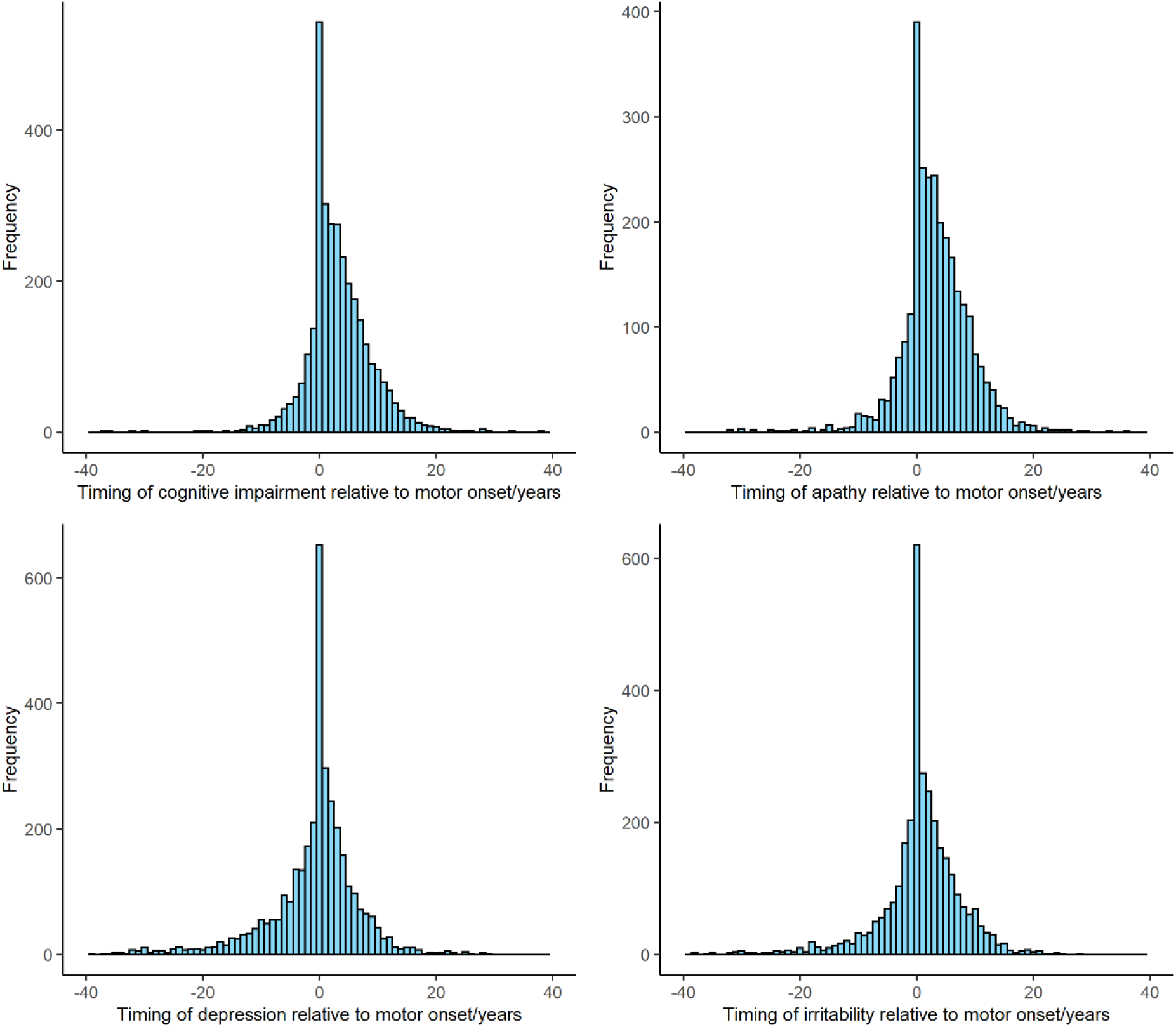
The onsets of cognitive and psychiatric symptoms relative to motor onset in HD. The age at onset of motor symptoms was subtracted from the age at onset of each cognitive/psychiatric symptom when present. Timings of up to +/− 40 years relative to motor onset shown. Only individuals with a rater-confirmed age at onset and CAG length (36-93) were included. Data from HD-CCQ. (**A**) Cognitive impairment N=3225; (**B**) Apathy N=2852; (**C**) Depression N=3495; (**D**) Irritability N=3235.

### The timing of motor and psychiatric symptoms in HD gene carriers varies with symptom type and CAG length

Given that motor onset is often used as a specific milestone in the natural history of HD, we investigated the timing of each of the seven psychiatric/cognitive symptoms relative to the age at first motor symptoms recorded in HD-CCQ (figure 2). The differences in ages between first motor symptoms and each of the psychiatric symptoms were approximately normally distributed, with a wide range of at least +/− 20 years in each case (figures 2 and e-2, doi:10.5061/dryad.pk0p2ngkz). In those patients reporting depression, onset occurred before motor symptoms in 39.2% (N= 1369/3495). For patients with irritability, onset occurred before motor symptoms in 30.8% (N= 996/3235). Perseverative/obsessive behaviour tended to occur later in the disease course, after motor symptoms, as did psychosis although numbers were smaller. Cognitive impairment and apathy had the most positively skewed distributions with onset occurring after motor onset in 2179/3225 (67.6%) and 1981/2852 (69.5%) of individuals, respectively. Overall, 42.4% of HD patients (N= 2140/5042) reported at least one psychiatric or cognitive symptom in advance of motor symptoms, with a further 22.3% (N= 1126/5042) reporting at least one of these symptoms at the same time as motor abnormalities.

We next assessed whether there were any patterns in the mean ages of onset of the different symptoms when plotted by CAG repeat length (figure 3). Some consistent relationships between symptoms were observed. Depression usually had the youngest mean age at onset, followed by motor impairment and then apathy and cognitive impairment as the latest symptoms. Mean age at onset of irritability preceded that of motor onset at shorter repeat lengths (40-43 CAGs, inclusive) but tended to follow it at longer repeat lengths (44-53 CAGs, inclusive). The mean difference in years from onset of first symptom to last decreased with CAG repeat length from approximately 8 years for 40 repeats to 4 years for 55 repeats (figure 3).

**Figure 3.**
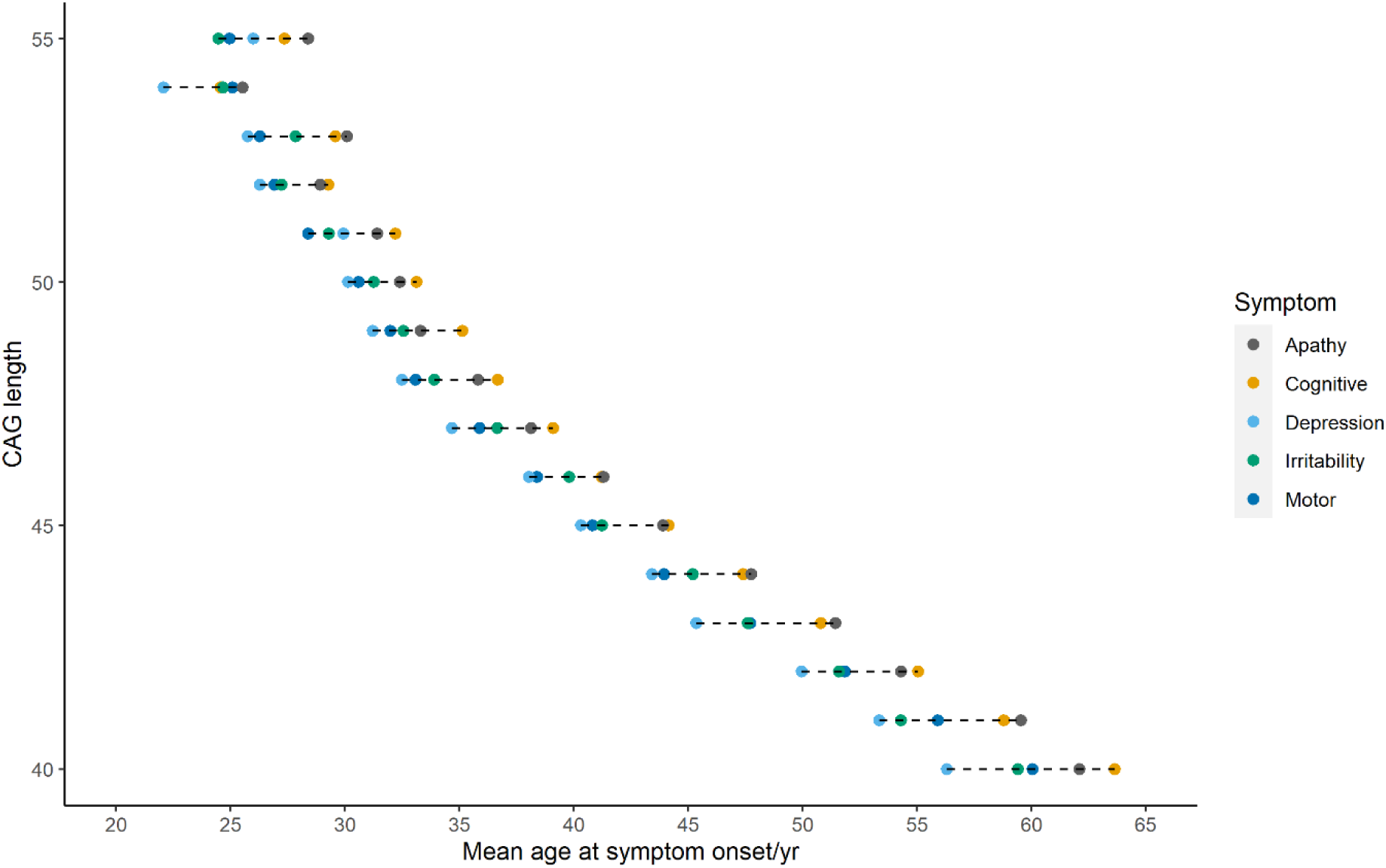
Mean ages at onset for motor and psychiatric symptoms at different CAG repeat lengths. Shown are the mean ages at symptom onset as recorded by the HD Clinical Characteristics Questionnaire for apathy (N = 2739), cognitive impairment (N = 3069), depression (N = 3399), irritability (N = 3117) and motor symptoms (N = 4889).

### Cognitive and psychiatric symptoms are significantly associated with reduced functional capacity

To assess whether psychiatric, cognitive or motor symptoms were associated with altered functional abilities we used multiple logistic regression (table 4). This analysis incorporated sex, pathogenic CAG length, duration of disease from clinical onset to last clinic visit, alcohol consumption, tobacco use, educational attainment, total functional capacity score and total motor score as predictors of the presence/absence of each HD-CCQ symptom. The presence of any of the psychiatric or cognitive symptoms was significantly associated with lower total functional capacity (TFC), an indication of impaired ability to work, manage personal finances and function independently. Cognitive impairment was most significantly associated with reduced TFC (OR per unit decrease in TFC = 1.28, 95% CI 1.23-1.35; *P* = 1.6 x 10^-25^). Depression was significantly associated with lower total motor scores (indicating fewer motor symptoms or signs), fitting with its prevalence early in the disease course. Finally, significant associations were observed between depression and female sex (OR = 1.77, 95% CI 1.44-2.17; *P* = 7.0 x 10^-8^) and tobacco use and irritability (OR per extra cigarette per day = 1.02, 95% CI 1.01-1.03; *P* = 1.0 x 10^-4^). Although not reaching strict criteria for significance after correcting for multiple tests, associations were also found between male sex and irritability (OR = 0.75, 95% CI 0.61-0.92; *P* = 5.4 x 10^-3^) and lower educational attainment and psychosis (OR per extra year of education = 0.92, 95% CI 0.88-0.97; *P* = 2.2 x 10^-3^).

## Discussion

In this large study of over 6000 patients we have shown that the initial manifestation of HD, as determined retrospectively by an expert rater, varies significantly with age. Late presentations (>60 years) are usually associated with motor abnormalities, whereas early presentations (<20 years; juvenile HD) are associated with a wider range of motor, cognitive and psychiatric abnormalities (figure 1A). These results extend prior studies which have shown that motor presentation of HD is common in late-onset disease (65.5% of an earlier REGISTRY cohort^24^), with more variable presentations in juvenile HD^25, 26^. Approximately 20% of HD patients present with rater-determined psychiatric features, in line with previous findings (table e-1, doi:10.5061/dryad.pk0p2ngkz)^9^. Cognitive onset of HD might be under-reported in older age-groups due to it being regarded as coincident ‘age-related’ change. Importantly, our results show there is little relationship between pathogenic CAG repeat length and onset type in adult-onset HD (figure 1B), despite both being associated with age at clinical onset. These data fit a model in which age at clinical onset is driven primarily by CAG repeat length, but modified by environmental factors and variants at other genomic loci^14, 23, 27, 28^. The age and physiology of the brain at clinical onset subsequently determines the types of symptoms that become manifest.

The HD-CCQ captures quantitative information not available elsewhere on symptom prevalence and timing in the HD population. Prior to its introduction in REGISTRY, age at first motor symptoms was not routinely recorded for all HD patients. HD-CCQ provides particular insight into neuropsychiatric symptoms but is not designed to capture the subtle early motor or cognitive signs found in prospective studies^7, 8^. As it relies on retrospective reporting by patients and care partners the HD-CCQ is necessarily coarse, although the data it generates correlate well with more precise measures of depression, irritability and cognition (table 1). Cognitive impairment measured by SDMT or Stroop correlated most strongly with lifetime history of cognitive impairment in HD-CCQ, as expected, but also showed significant correlations with motor symptoms and apathy. These results fit with other studies showing that these symptoms track together in the disease trajectory^29, 30^. There was also a significant association between cognitive impairment and psychosis, which fits the cognitive deficits observed in schizophrenia^31^. Conversely, validated depression and irritability scores correlated well with their respective prevalence data from HD-CCQ but were not associated with motor or cognitive impairment (table 1).

Almost all HD patients have specific motor abnormalities consistent with HD during their disease course (table 2). Psychiatric and cognitive symptoms are also very common (table 2), much more prevalent than in non-HD populations^5, 10, 32, 33^, and likely underestimated due to pathological unawareness of these traits by HD patients^34^. However, clinically it is currently impossible to distinguish between symptoms arising as a result of the HD mutation and those from primary psychiatric disorders, particularly in younger pre-manifest patients in whom diseases such as depression are common^35^. Furthermore, environmental effects on mental health, such as living in a family with HD, should not be overlooked. As such, non-motor symptoms should not be used to make a clinical diagnosis of HD and this could even cause harm in vulnerable individuals with psychiatric symptoms. Future studies of psychiatric and cognitive symptoms and signs in HD gene carriers against gene-negative community controls might help define a HD-specific neuropsychiatric phenotype that would enable more confident attribution of early abnormalities to HD.

The age at onset of each symptom recorded by HD-CCQ was inversely correlated with CAG length (figure 3), with motor symptoms best correlated (table 3). Depression was least correlated (R^2^ = 37.5%), likely reflecting the high prevalence of the symptom in the general population independent of HD and the lack of use of universal diagnostic criteria. These data are consistent with previous reports showing that CAG length accounts for 47-72% of the variance in age at motor onset of HD^36^, but contradict previous studies that reported no correlation between CAG repeat length and psychiatric symptoms^37–40^. However, these studies were small and often examined incident psychiatric symptoms, which can fluctuate over time, rather than lifetime history as here. Increasingly accurate CAG tract sizing will improve the accuracy of correlations between repeat length and symptoms^14, 41, 42^.

**Table 3.** Variance in age at onset (R^2^) explained by pathogenic CAG repeat length for eight symptoms in males and females with HD. Ages at onset were logarithmically transformed and plotted against CAG length. *P* values test difference between male and female R^2^. Significant *P* values (*P* < 6.25 x 10^-3^; multiple testing correction) are in bold and nominally significant *P* values (*P* < 0.05) italicised. Individuals had to have a clinical onset of HD, a known sex and a pathogenic CAG length (36-93) to be included. CI: confidence interval; POB: perseverative/obsessive behaviour; VAB: violent or aggressive behaviour.

|  | Male |  | Female |  | *P* | Both |  |
| --- | --- | --- | --- | --- | --- | --- | --- |
|  | R<sup>2</sup> (95% CI) | N | R<sup>2</sup> (95% CI) | N |  | R<sup>2</sup> (95% CI) | N |
| Motor | 0.678<br>(0.657 - 0.697) | 2684 | 0.649<br>(0.628 - 0.670) | 2844 | 5.42 x 10<sup>-2</sup> | 0.663<br>(0.648 - 0.677) | 5528 |
| Cognitive | 0.610<br>(0.579 - 0.639) | 1570 | 0.629<br>(0.600 - 0.656) | 1681 | 3.80 x 10<sup>-1</sup> | 0.619<br>(0.598 - 0.639) | 3251 |
| Apathy | 0.595<br>(0.562 - 0.627) | 1423 | 0.562<br>(0.528 - 0.595) | 1462 | 1.83 x 10<sup>-1</sup> | 0.578<br>(0.554 - 0.601) | 2885 |
| Depression | 0.412<br>(0.374 - 0.449) | 1551 | 0.351<br>(0.318 - 0.385) | 1994 | 3.50 x 10<sup>-2</sup> | 0.375<br>(0.350 - 0.400) | 3545 |
| POB | 0.539<br>(0.496 - 0.581) | 973 | 0.440<br>(0.394 - 0.485) | 1016 | **3.67 x 10<sup>-3</sup>** | 0.489<br>(0.457 - 0.52) | 1989 |
| Irritability | 0.463<br>(0.428 - 0.498) | 1670 | 0.547<br>(0.513 - 0.579) | 1601 | **1.25 x 10<sup>-3</sup>** | 0.503<br>(0.478 - 0.527) | 3271 |
| VAB | 0.479<br>(0.431 - 0.524) | 927 | 0.478<br>(0.426 - 0.528) | 761 | 9.79 x 10<sup>-1</sup> | 0.477<br>(0.442 - 0.511) | 1688 |
| Psychosis | 0.401<br>(0.316 - 0.484) | 312 | 0.424<br>(0.340 - 0.504) | 318 | 7.29 x 10<sup>-1</sup> | 0.411<br>(0.351 - 0.469) | 630 |

Despite considerable variation in the timing of psychiatric and motor symptoms, there are some conserved patterns (figures 2 & 3). Depression, and less often irritability, can precede motor symptoms by many years. Conversely, apathy and cognitive impairment tend to occur after motor symptoms, although patients do recognise and report these symptoms less readily than depression or irritability. Overall, the HD-CCQ data show that 64.8% of our HD population (N = 3266/5042) reported at least one psychiatric or cognitive symptom by the time of first motor symptoms. This is a much higher figure than previously reported based on clinician estimates of first HD manifestation (figure 1)^9^, most likely because it is difficult to confidently attribute early psychiatric symptoms to HD. The overlap between HD and psychiatric disorders has been demonstrated by the recent finding that polygenic risk scores for psychiatric diseases, particularly depression and schizophrenia, are associated with increased risk of corresponding psychiatric symptoms in HD^29^. This suggests that the expanded *HTT* CAG repeat might lower the genetic threshold for manifestation of typical psychiatric symptoms^29^. In agreement, we found the expected relationships between female sex and depression and male sex and irritability in our cohort (table 4). The nominally significant negative association of psychosis in HD with educational level (table 4) also corroborates work showing that higher levels of education are associated with decreased schizophrenia risk^43^.

**Table 4.**
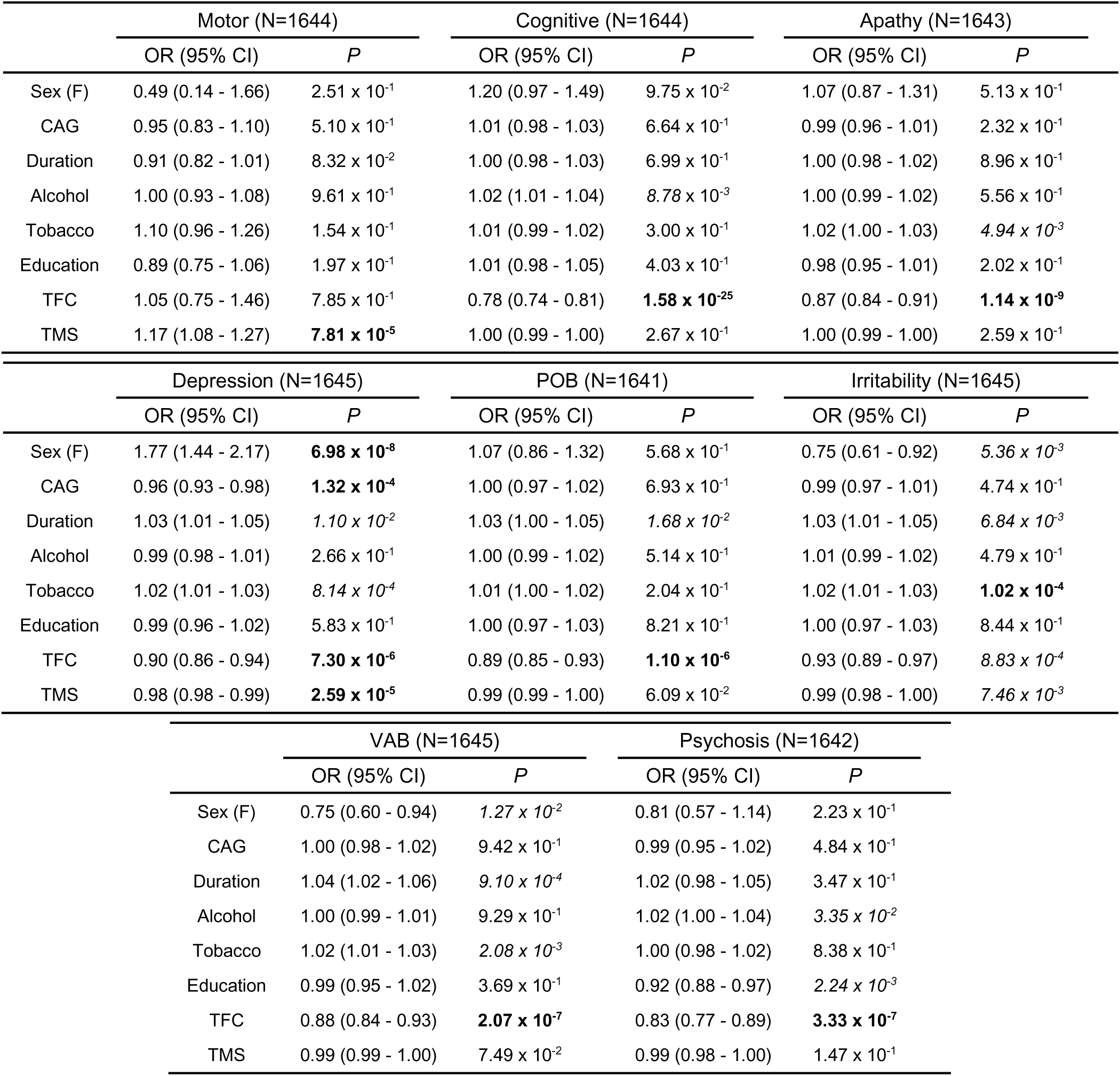
Psychiatric and cognitive symptoms are associated with reduced functional capacity. Multiple logistic regression using binary HD-CCQ data for 8 symptoms (0 = no symptom; 1 = reported symptom) and clinical covariates. Significant associations after Bonferroni correction for 8 symptoms and 8 covariates are shown in bold (*P* < 7.81 x 10^-4^) and nominally significant associations in italics (*P* < 0.05). With the exception of sex, the odds ratio (OR) indicates the effect on the outcome probability associated with an increase of one unit in the covariate. In addition to having a confirmed onset and pathogenic CAG length (36-93), individuals must have no co-morbid diagnosis of schizophrenia, schizotypy or schizoaffective disorder. CI: confidence interval; Sex (F): female; POB: perseverative/obsessive behaviour; TFC: total functional capacity; TMS: total motor score; VAB: violent or aggressive behaviour.

We acknowledge several potential limitations of these data: they are retrospective, subject to recall bias, and cross-sectional. Furthermore, HD-CCQ data depend on the interpretation of questions: for example, ‘motor symptoms’ are not explicitly defined and so although 96.8% of our population had chorea this was not documented in HD-CCQ. Future iterations might usefully subdivide ‘motor symptoms’ into (1) fidgety or jerky involuntary movements (chorea)’ and (2) other HD-related movement problems such as unsteadiness, stiffness or trouble with fine movements. Our analyses are based on data from the most recent clinic visit which is at different points of the disease course in different individuals. We controlled for this by using disease duration, the time between first onset and last clinic visit, as a covariate in analyses. The use of psychoactive medications is found in up to 60% of HD patients and might confound motor and neuropsychiatric phenotypes^9, 44^. Of drugs prescribed for chorea, tetrabenazine can induce depression and antipsychotics can reduce irritability. They also suppress motor manifestations which might affect the total motor scores used here as a co-variate (table 4). It is hard to control for these effects. Drugs prescribed to treat symptoms once they are present will not influence symptom onset data. We used worst-ever depression and irritability scores when validating the use of HD-CCQ to mitigate against the effects of medication prescribed at certain times.

Previous prospective studies of phenotype in HD such as PREDICT-HD and TRACK-HD have shown subtle early reductions in psychiatric and cognitive function years in advance of clinical onset^7, 8^. The HD-CCQ accesses retrospective data from large existing populations of manifest HD patients and shows similar trends. Since the HD-CCQ is part of ongoing longitudinal observational studies such as ENROLL-HD, future analyses of larger populations will be possible and of benefit. The presence of psychiatric and cognitive symptoms in HD gene carriers is associated with significantly reduced functional capacity, emphasizing the importance of early recognition and management of these symptoms^45, 46^. Although recent models of HD staging and progression do not directly include psychiatric and cognitive symptoms^47–49^, work is underway to include them in ongoing observational studies and clinical trials to improve the accuracy of clinical outcome measures.

## Acknowledgments

We thank all the patients who contributed data to this research. B.Mc. was supported by a PhD studentship from Cardiff University School of Medicine. J.F.G. and M.E.M. received support from NIH grant NS091161 and from the CHDI Foundation, Inc. J-M.L. received support from grant R01NE-105709. A.E.R. received support from MRC, Wellcome Trust, Campaign for Alzheimer’s Research in Europe, Horizon 2020, JPND and Health and Care Research Wales. L.J., N.M.W. and P.H. were supported by an MRC Centre grant (MR/L010305/1). T.H.M. was supported by a Welsh Clinical Academic Track Fellowship, an MRC Clinical Training Fellowship (MR/P001629/1) and a Patrick Berthoud Charitable Trust Fellowship through the Association of British Neurologists.

## Financial Disclosures of all authors (for the preceding 12 months)

J.F.G.: Scientific Advisory Board member and has a financial interest in Triplet Therapeutics, Inc. His NIH-funded project is using genetic and genomic approaches to uncover other genes that significantly influence when diagnosable symptoms emerge and how rapidly they worsen in Huntington Disease. The company is developing new therapeutic approaches to address triplet repeat disorders such Huntington’s Disease, Myotonic Dystrophy and spinocerebellar ataxias. His interests were reviewed and are managed by Massachusetts General Hospital and Partners HealthCare in accordance with their conflict of interest policies.

G.B.L.: Consulting services, advisory board functions, clinical trial services and/or lectures for Allergan, Alnylam, Amarin, AOP Orphan Pharmaceuticals AG, Bayer Pharma AG, CHDI Foundation, GlaxoSmithKline, Hoffmann-LaRoche, Ipsen, ISIS Pharma, Lundbeck, Neurosearch Inc, Medesis, Medivation, Medtronic, NeuraMetrix, Novartis, Pfizer, Prana Biotechnology, Sangamo/Shire, Siena Biotech, Temmler Pharma GmbH and Teva Pharmaceuticals. He has received research grant support from the CHDI Foundation, the Bundesministerium für Bildung und Forschung (BMBF), the Deutsche Forschungsgemeinschaft (DFG), the European Commission (EU-FP7, JPND). His study site Ulm has received compensation in the context of the observational Enroll-HD Study, TEVA, ISIS and Hoffmann-Roche and the Gossweiler Foundation. He receives royalties from the Oxford University Press and is employed by the State of Baden-Württemberg at the University of Ulm.

A.E.R.: Chair of European Huntington’s Disease Network (EHDN) executive committee, Global PI for Triplet Therapeutics

L.J. is a member of the scientific advisory boards of LoQus23 Therapeutics and Triplet Therapeutics and has received grant support from CHDI.

T.H.M. is an associate member of the scientific advisory board of LoQus23 Therapeutics.

B.Mc., J-M.L., M.E.M., M.O., N.M.W., P.H.: nothing to disclose

## Appendix 1 Authors

**Appendix 1 – Authors**
| Name | Location | Contribution/Role |
| --- | --- | --- |
| Branduff McAllister, BSc, PhD | Cardiff University, Cardiff, UK | Organized data; Designed and executed statistical analyses; Wrote first paper draft; Reviewed and critiqued paper manuscript |
| James F. Gusella, PhD | Massachusetts General Hospital, Boston, USA | Reviewed and critiqued paper manuscript |
| Bernhard Landwehrmeyer, MD, PhD | University of Ulm, Ulm, Germany | Reviewed and critiqued paper manuscript |
| Jong-Min Lee, PhD | Massachusetts General Hospital, Boston, USA | Reviewed and critiqued paper manuscript |
| Marcy E. MacDonald, PhD | Massachusetts General Hospital, Boston, USA | Reviewed and critiqued paper manuscript |
| Michael Orth, MD, PhD | Swiss Huntington's disease Centre, Bern, Switzerland | Reviewed and critiqued paper manuscript |
| Anne E. Rosser, MB BChir, FRCP, PhD | Cardiff University, Cardiff, UK | Reviewed and critiqued paper manuscript |
| Nigel M. Williams, BSc, PhD | Cardiff University, Cardiff, UK | Reviewed and critiqued paper manuscript |
| Peter Holmans, BA, PhD | Cardiff University, Cardiff, UK | Designed and conceptualized study; Designed and critiqued statistical analyses; Reviewed and critiqued paper manuscript |
| Lesley Jones, BSc, PhD (*) | Cardiff University, Cardiff, UK | Designed and conceptualized study; Wrote first paper draft; Reviewed and critiqued paper manuscript |
| Thomas H. Massey, MA, BM Bch, DPhil (*), (†) | Cardiff University, Cardiff, UK | Designed and conceptualized study; Wrote first paper draft; Reviewed and critiqued paper manuscript |

2 (\*) indicates corresponding authors
3 (†) indicates the principal investigator

## Appendix 2 Co-investigators of the European Huntington’s Disease Network REGISTRY Study

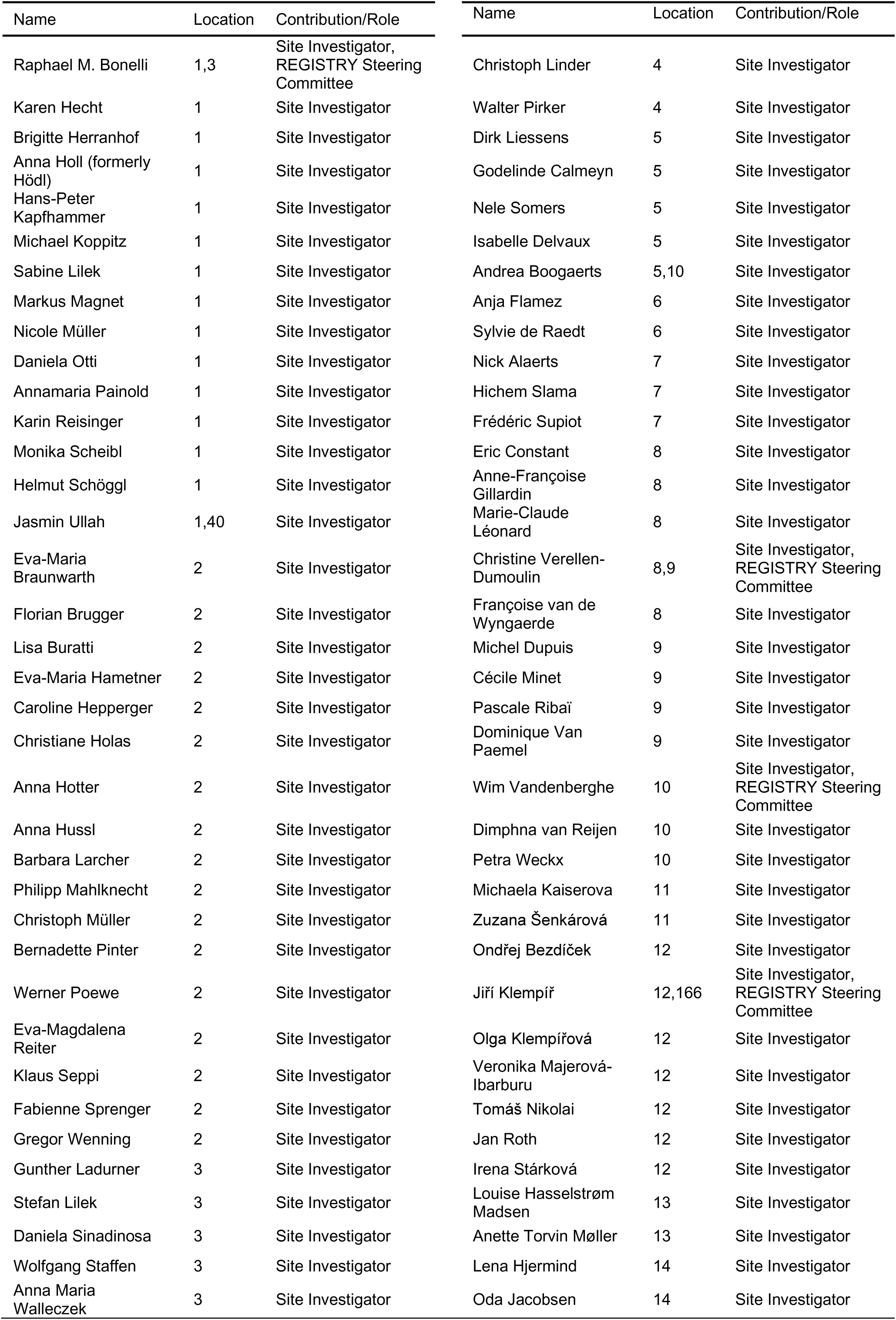
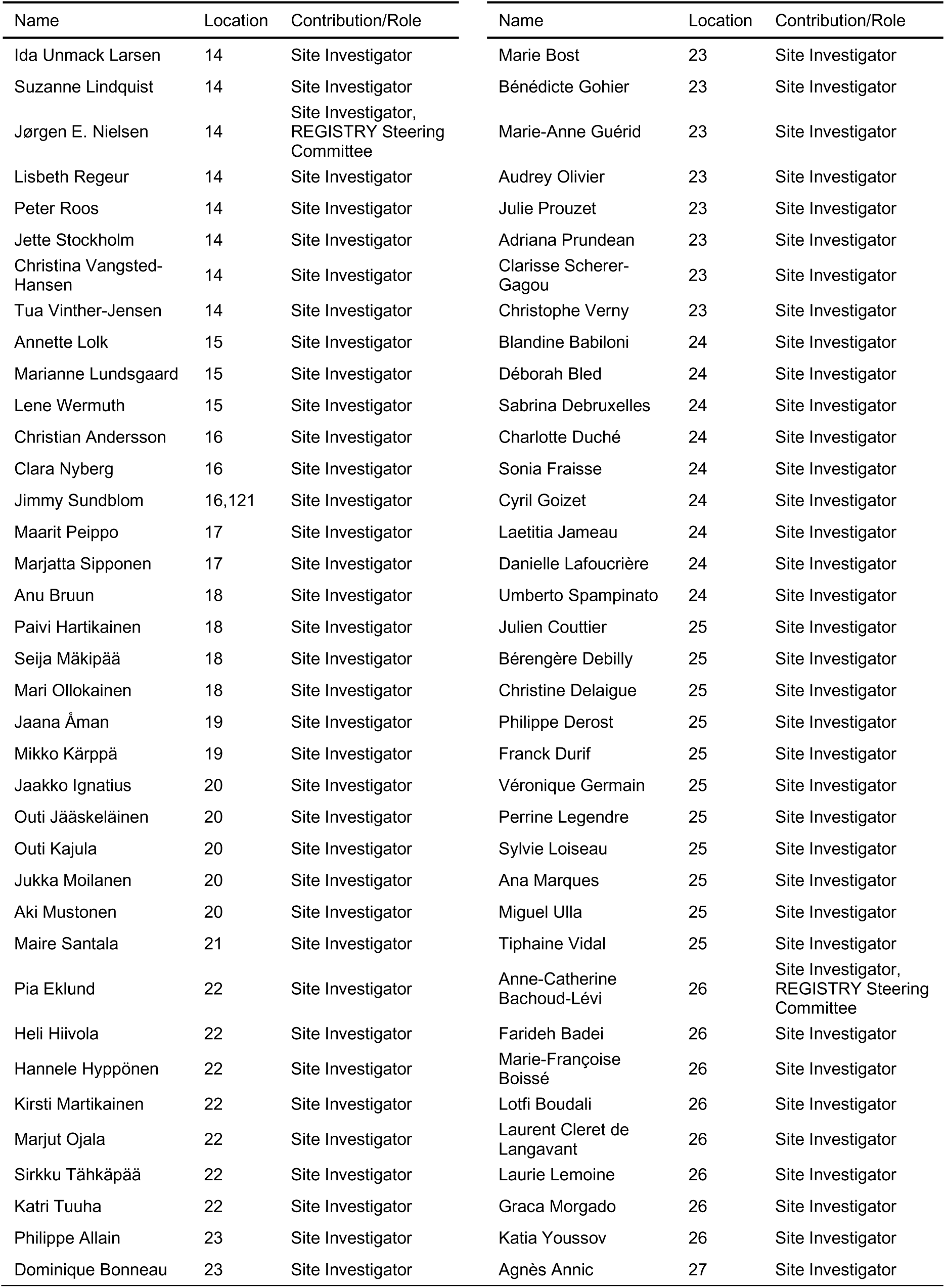
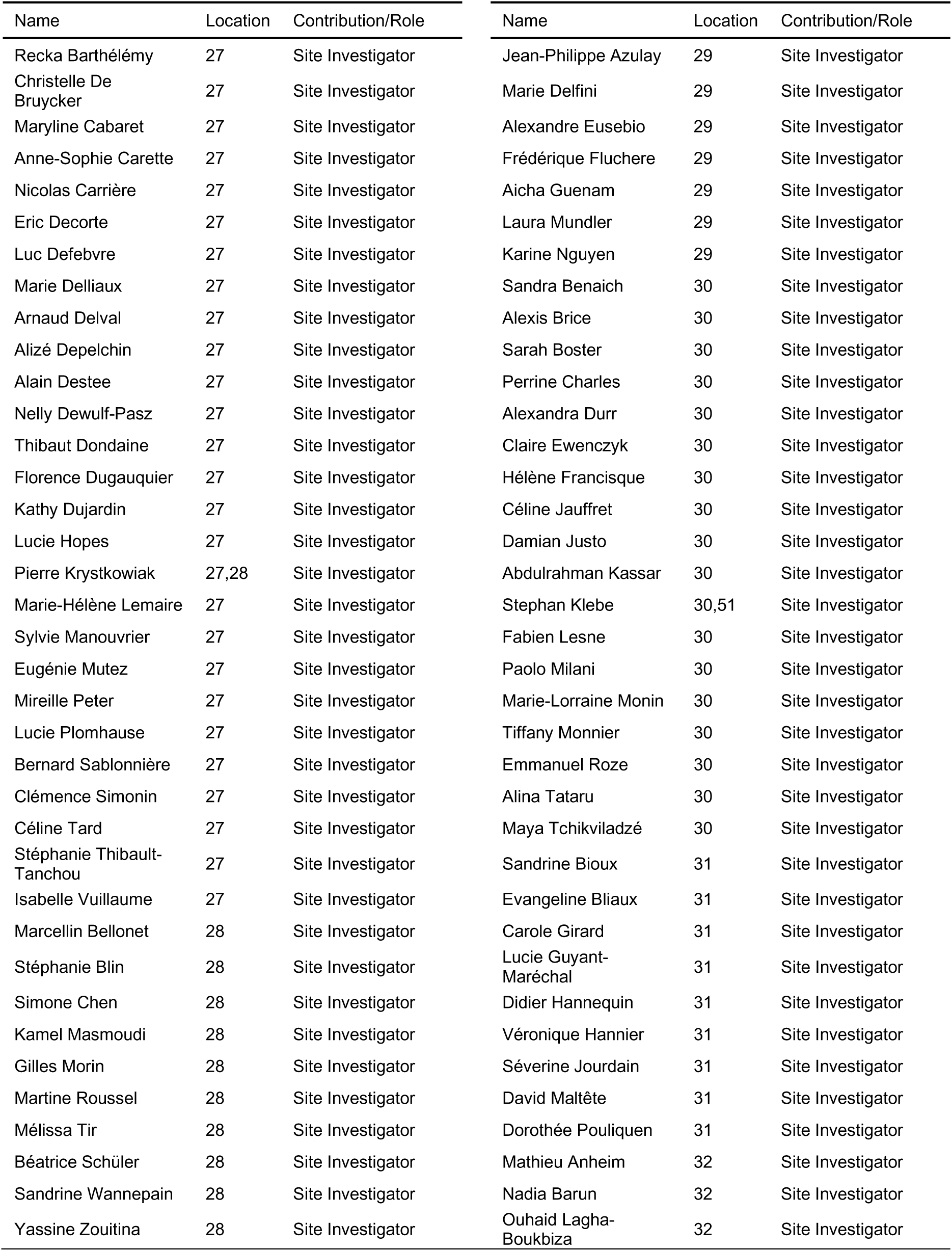
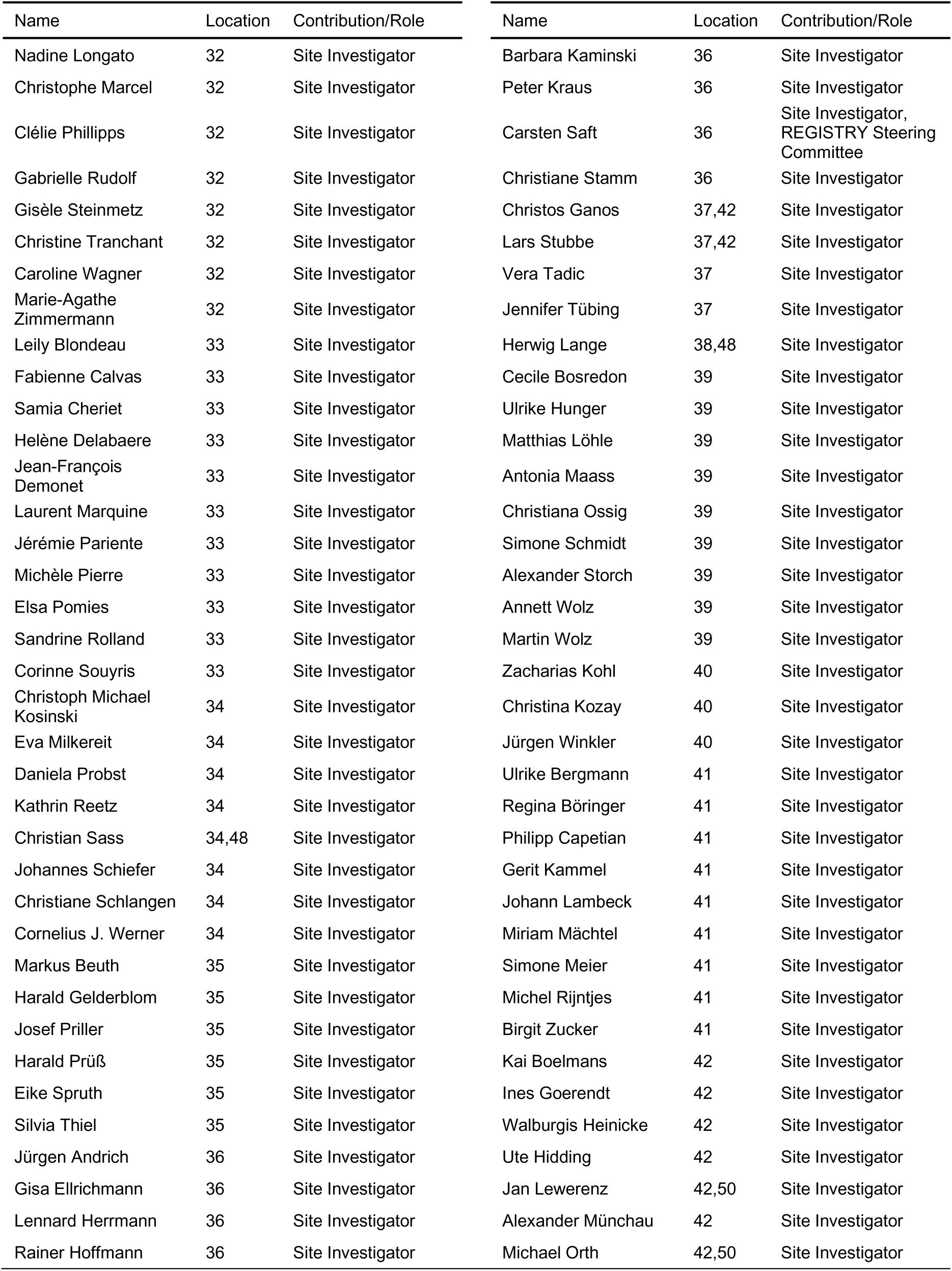
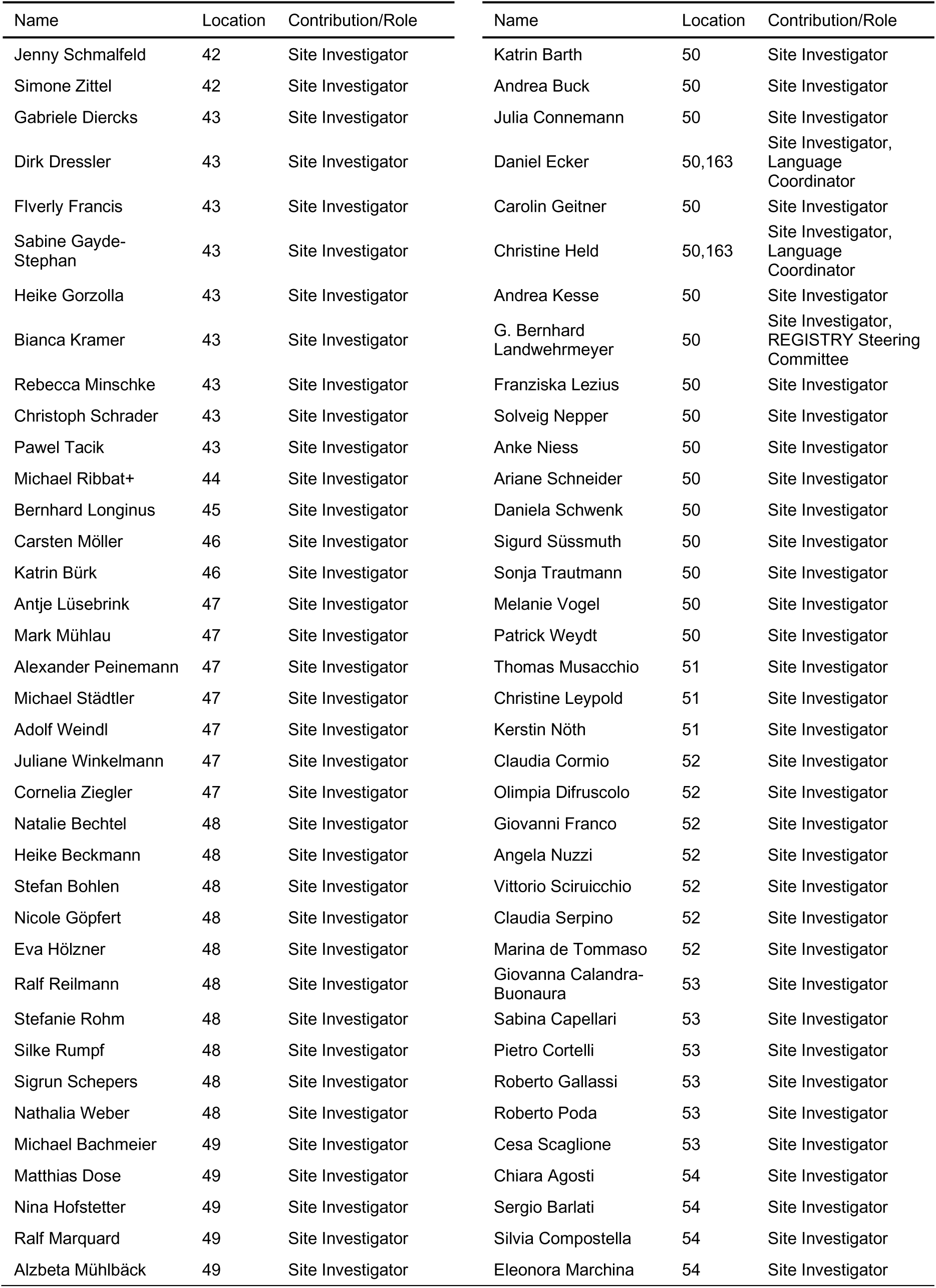
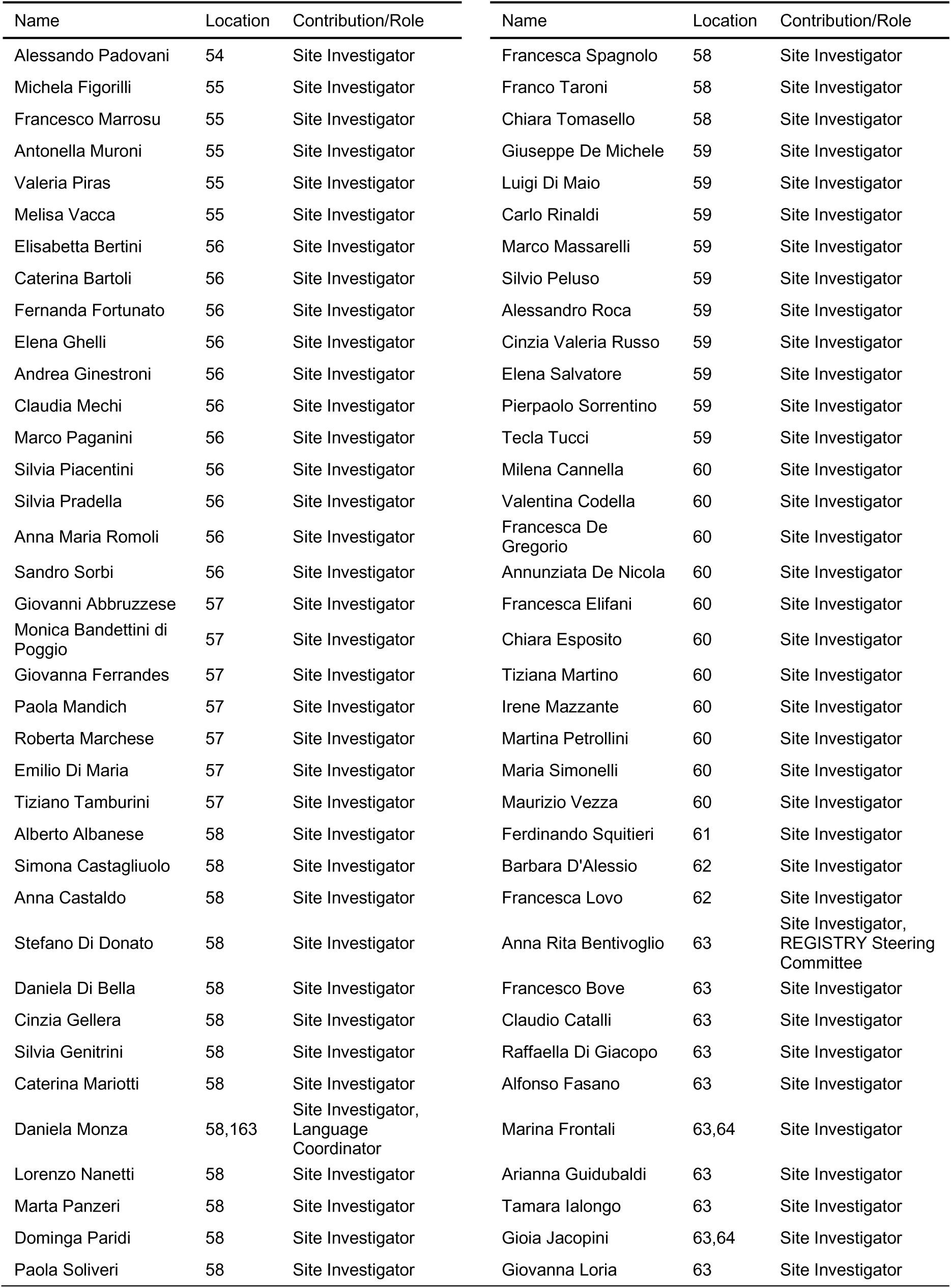
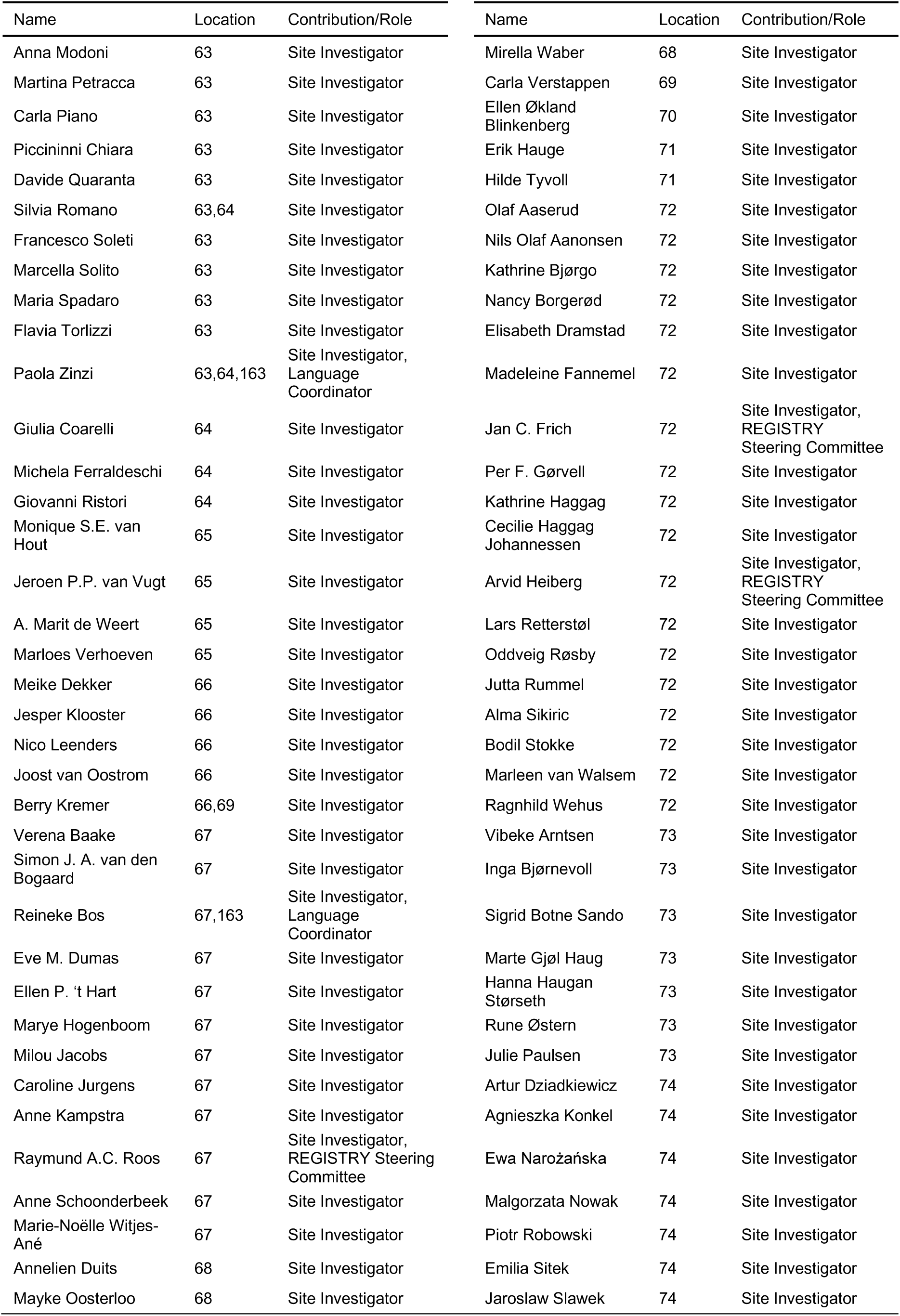
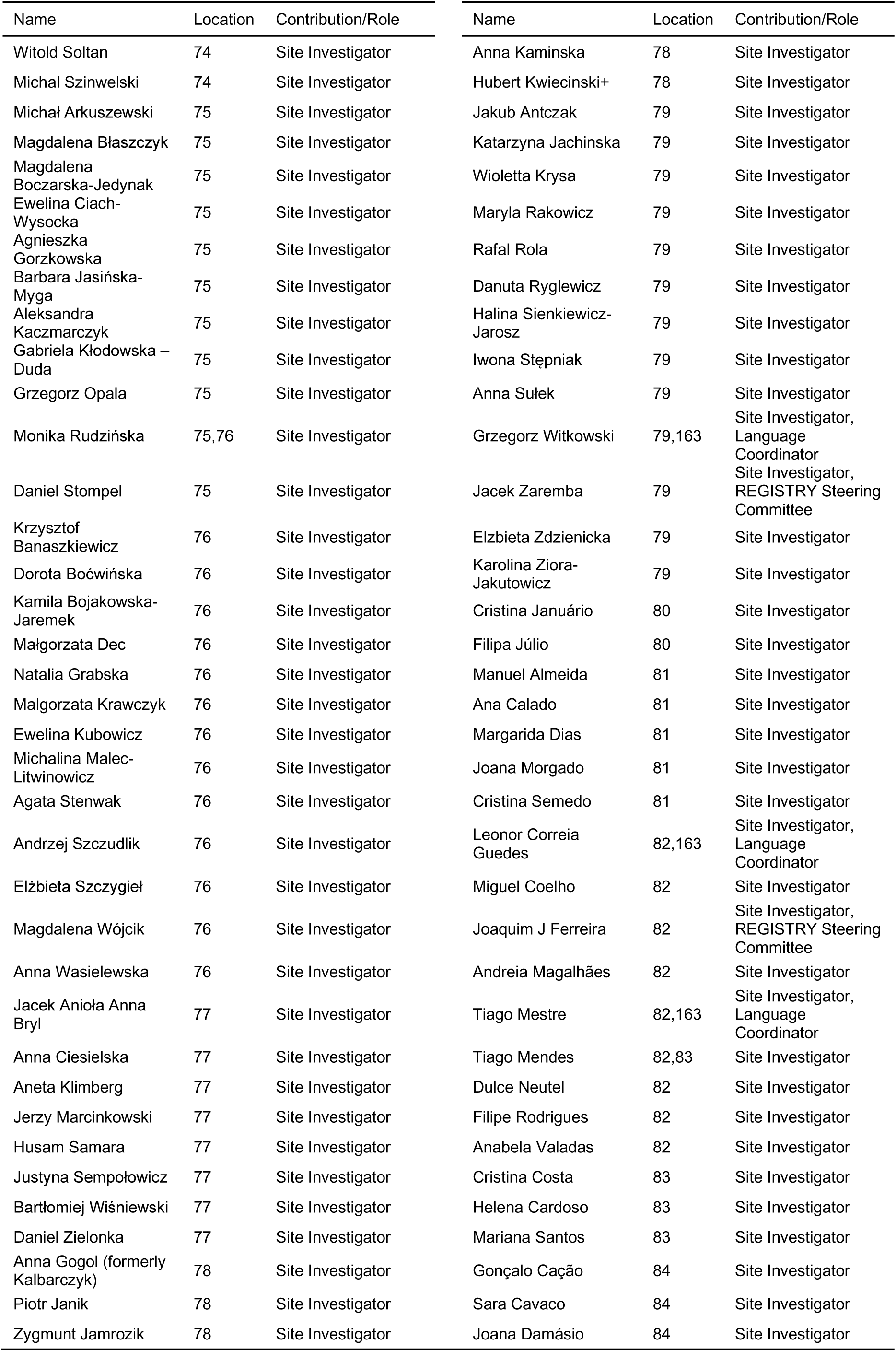
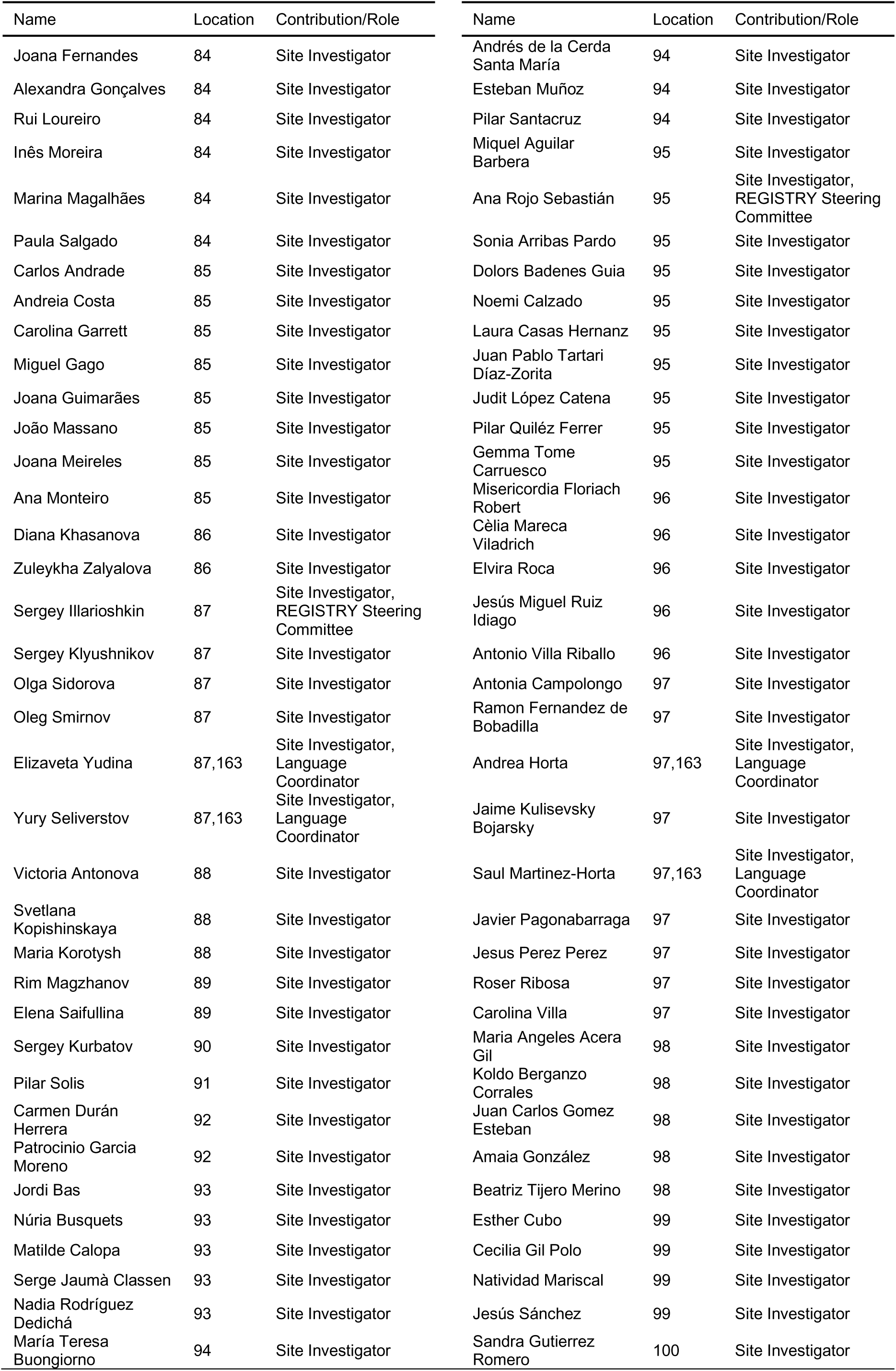
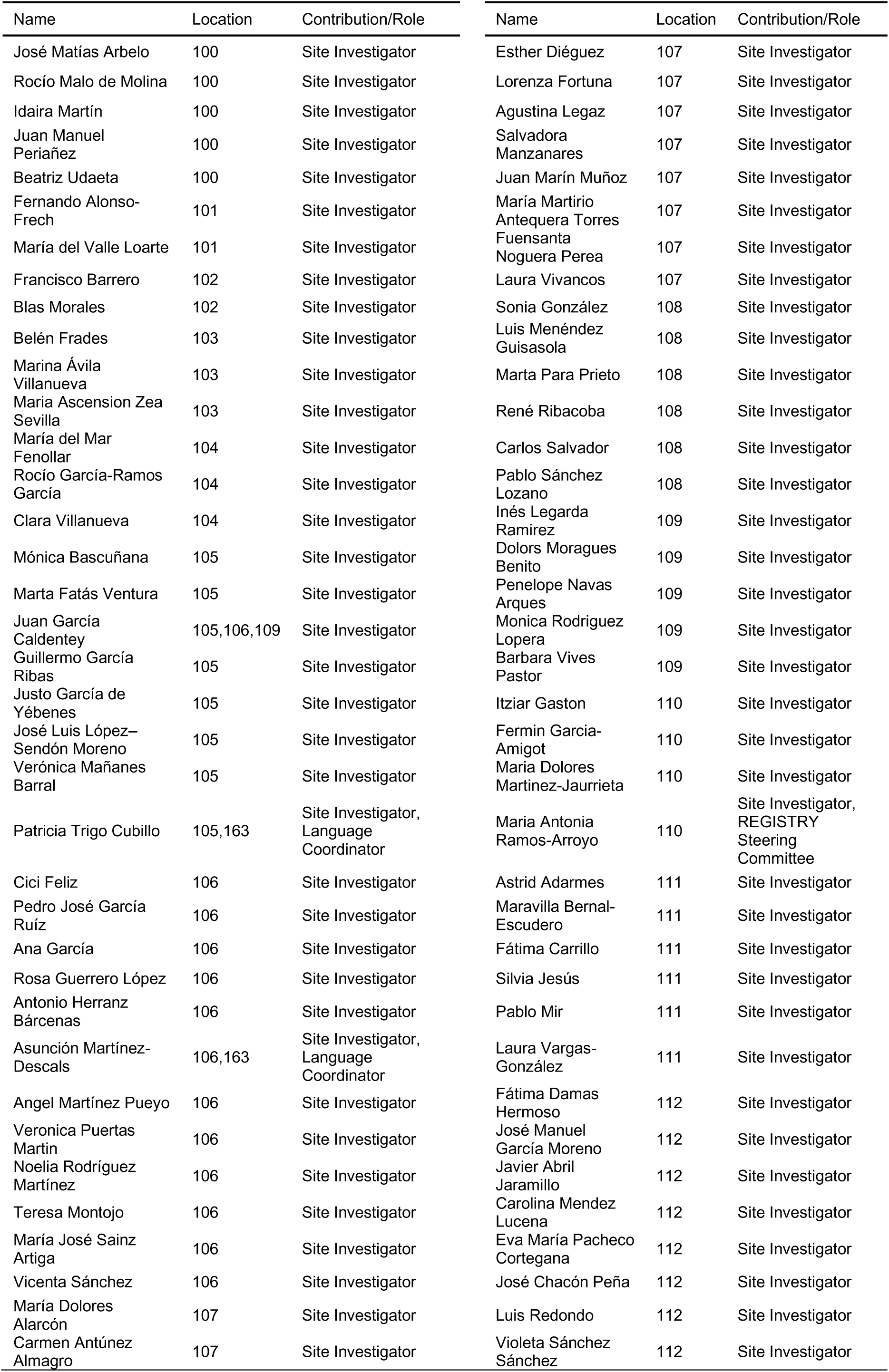
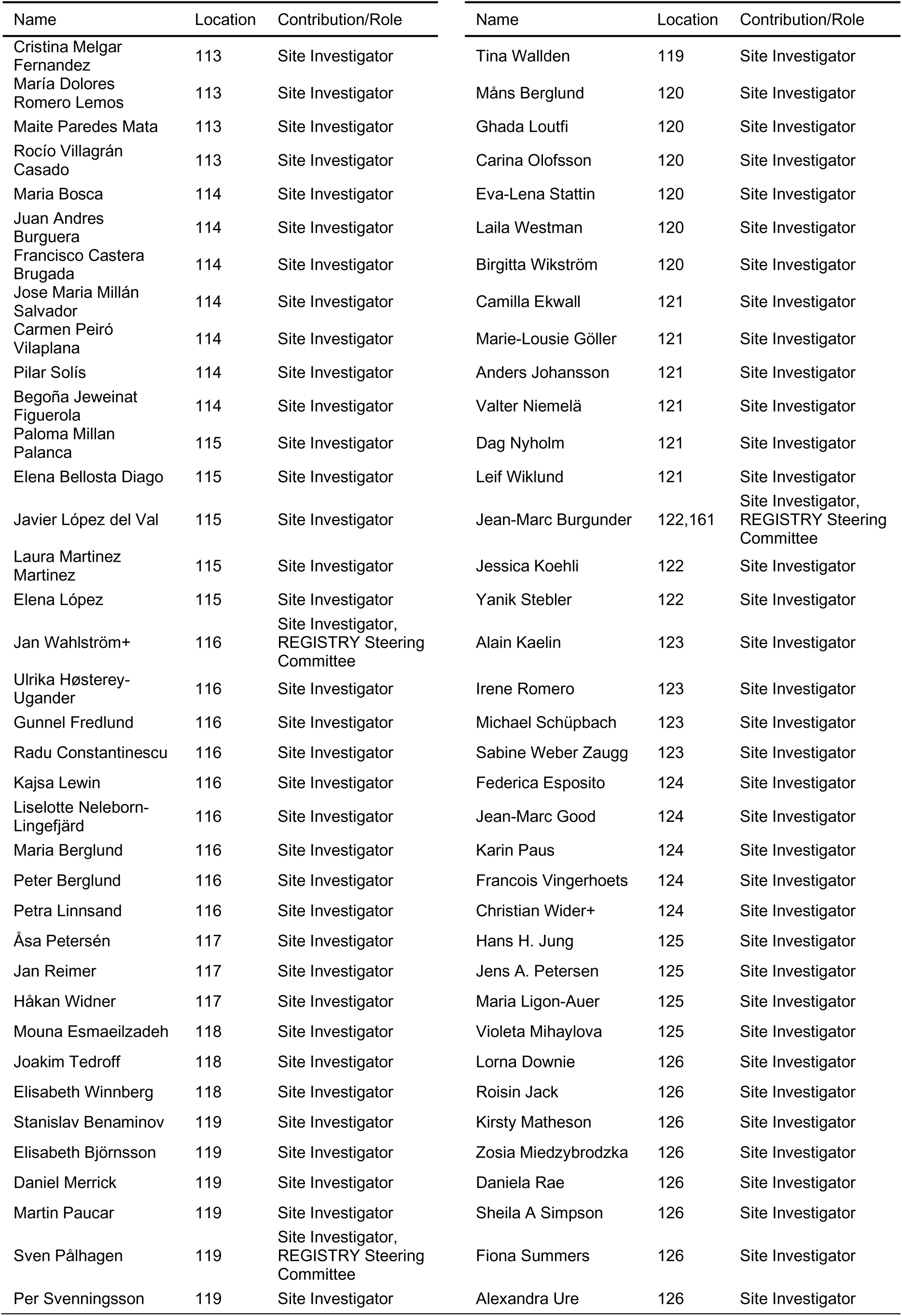
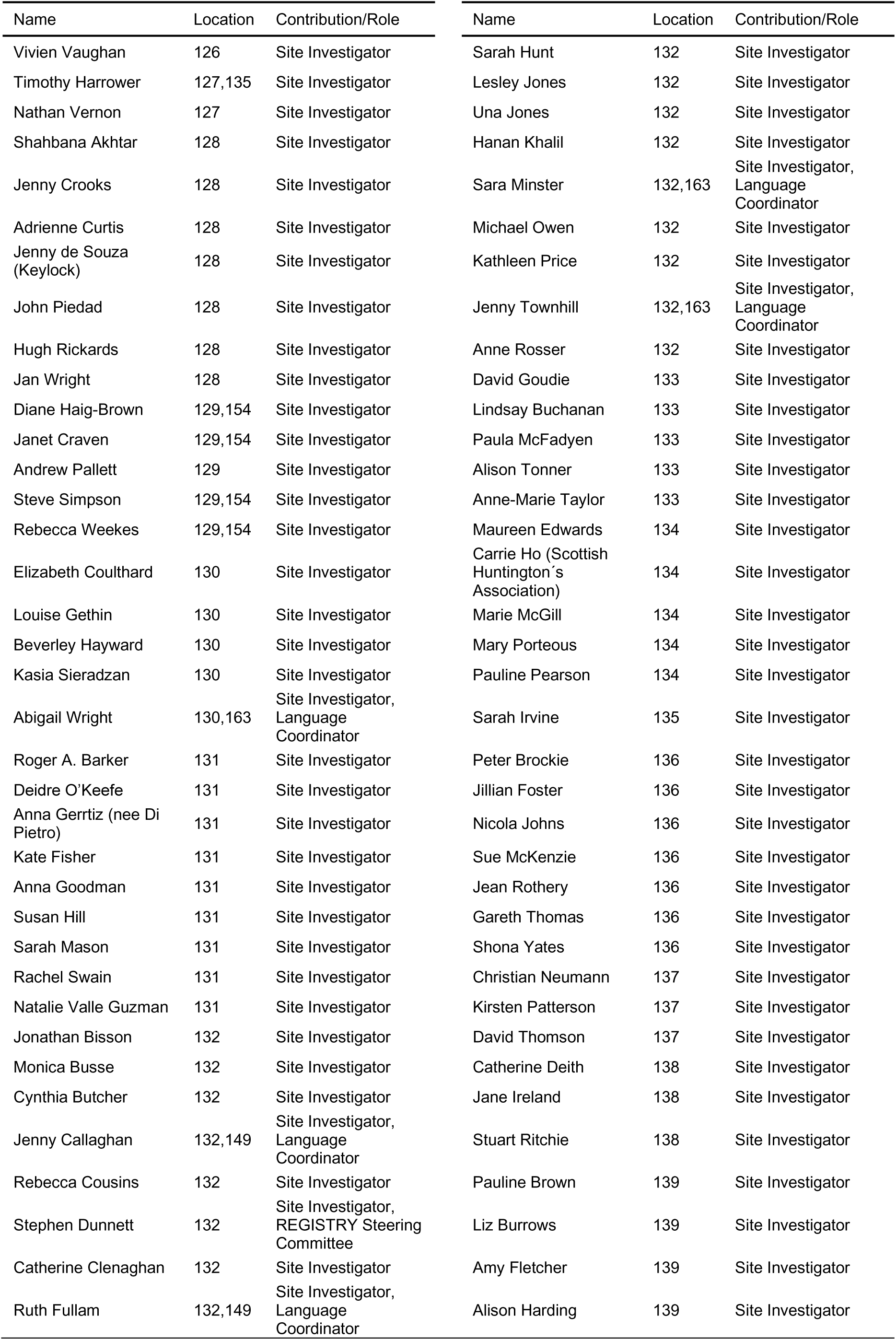
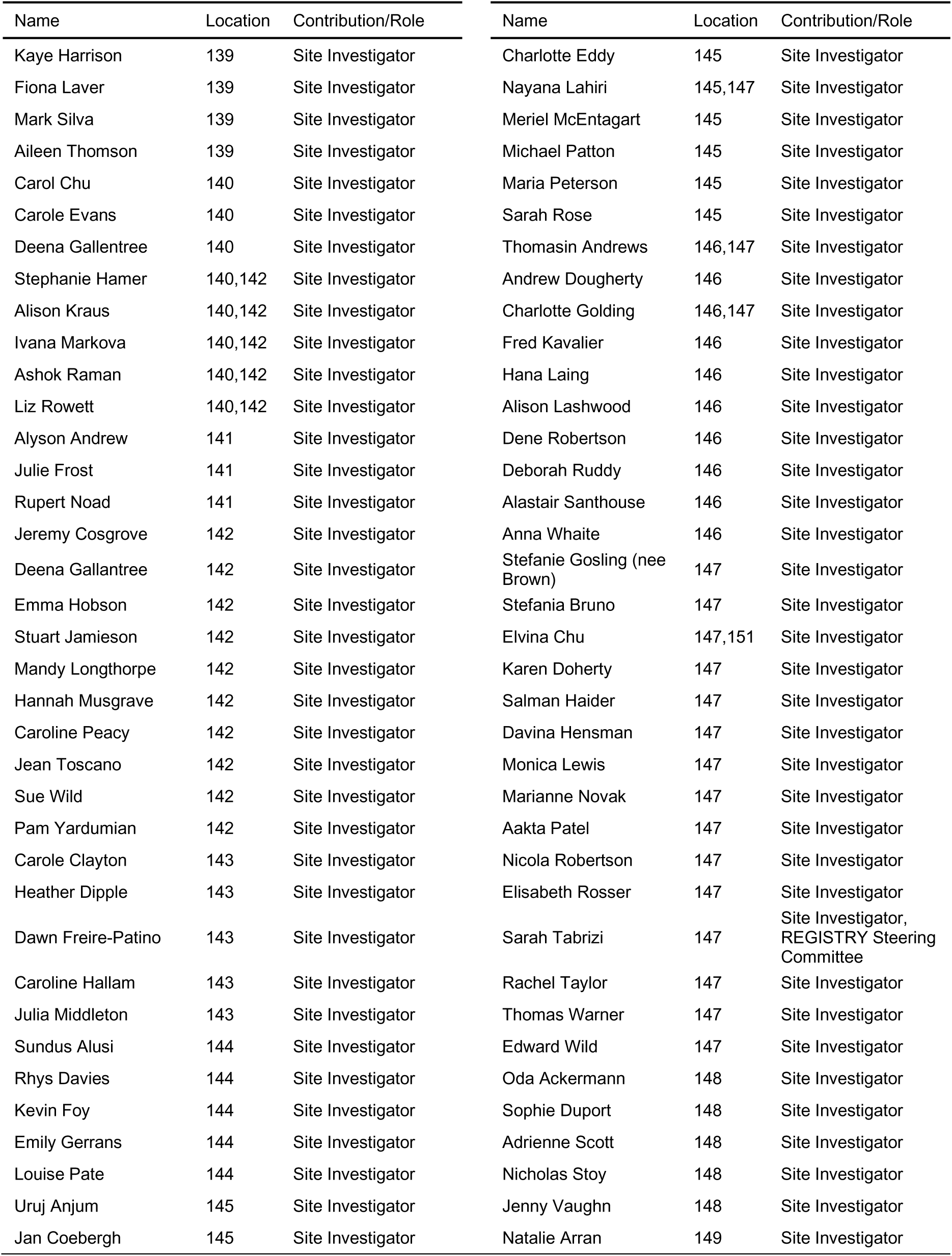
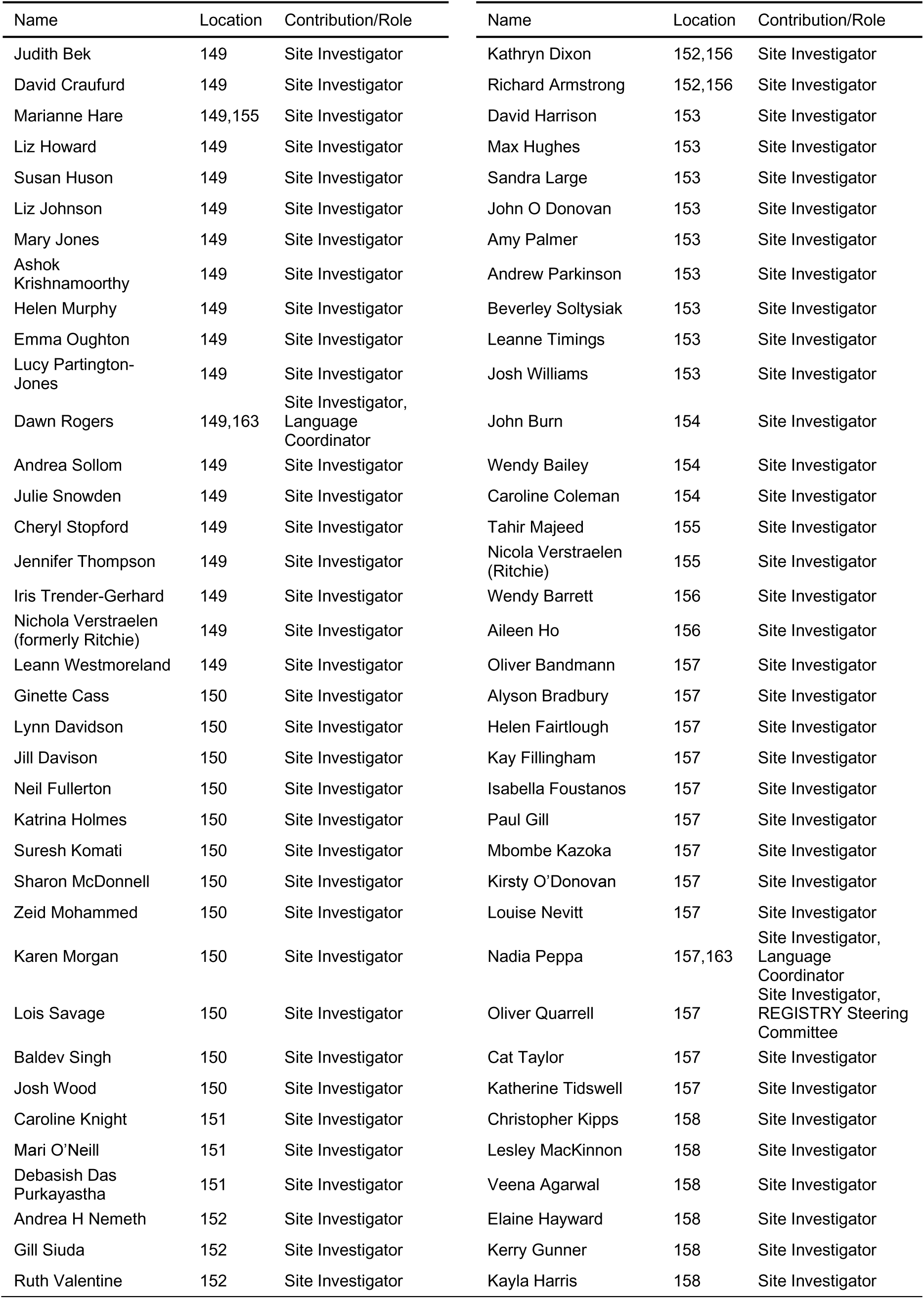
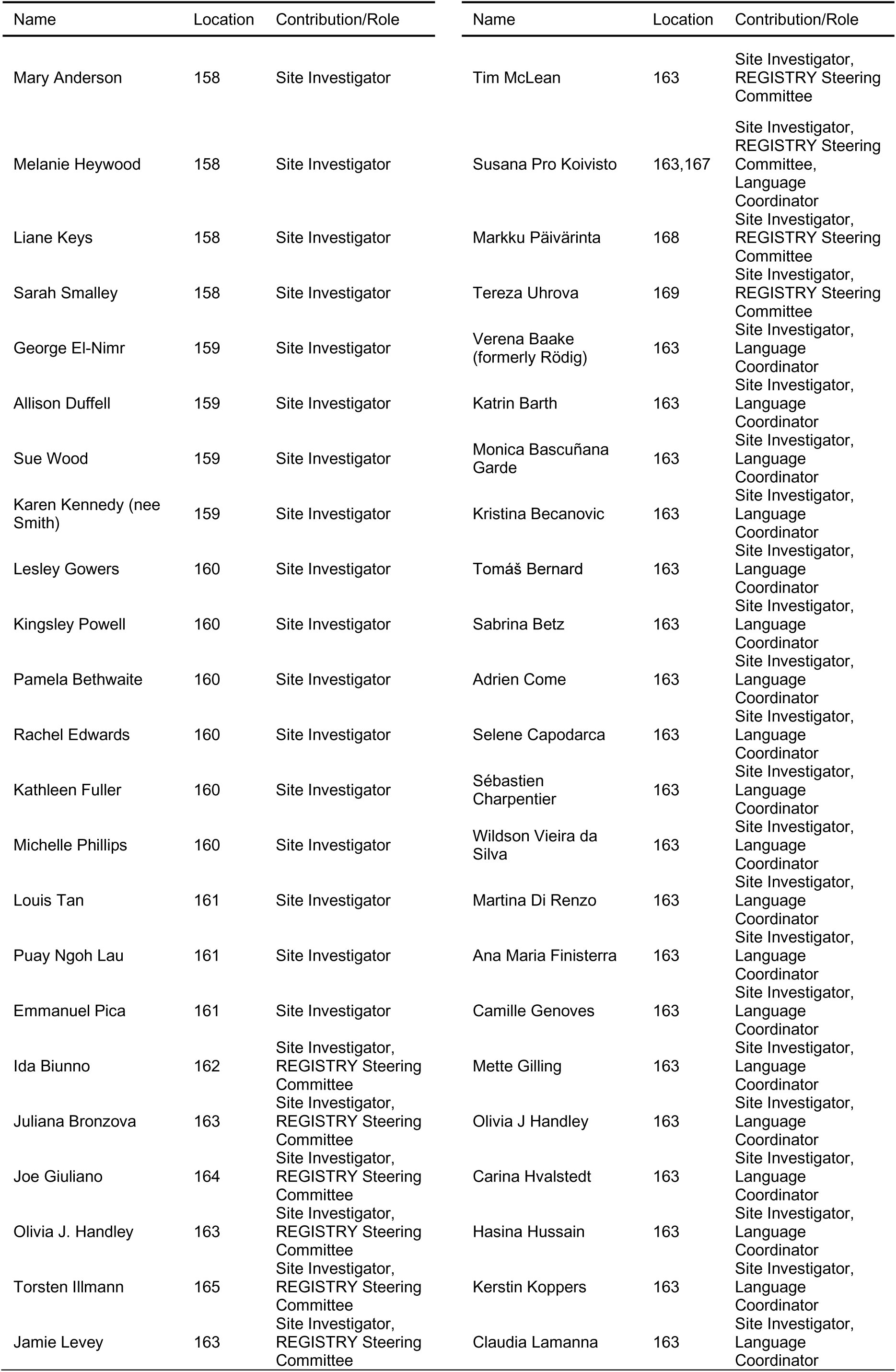
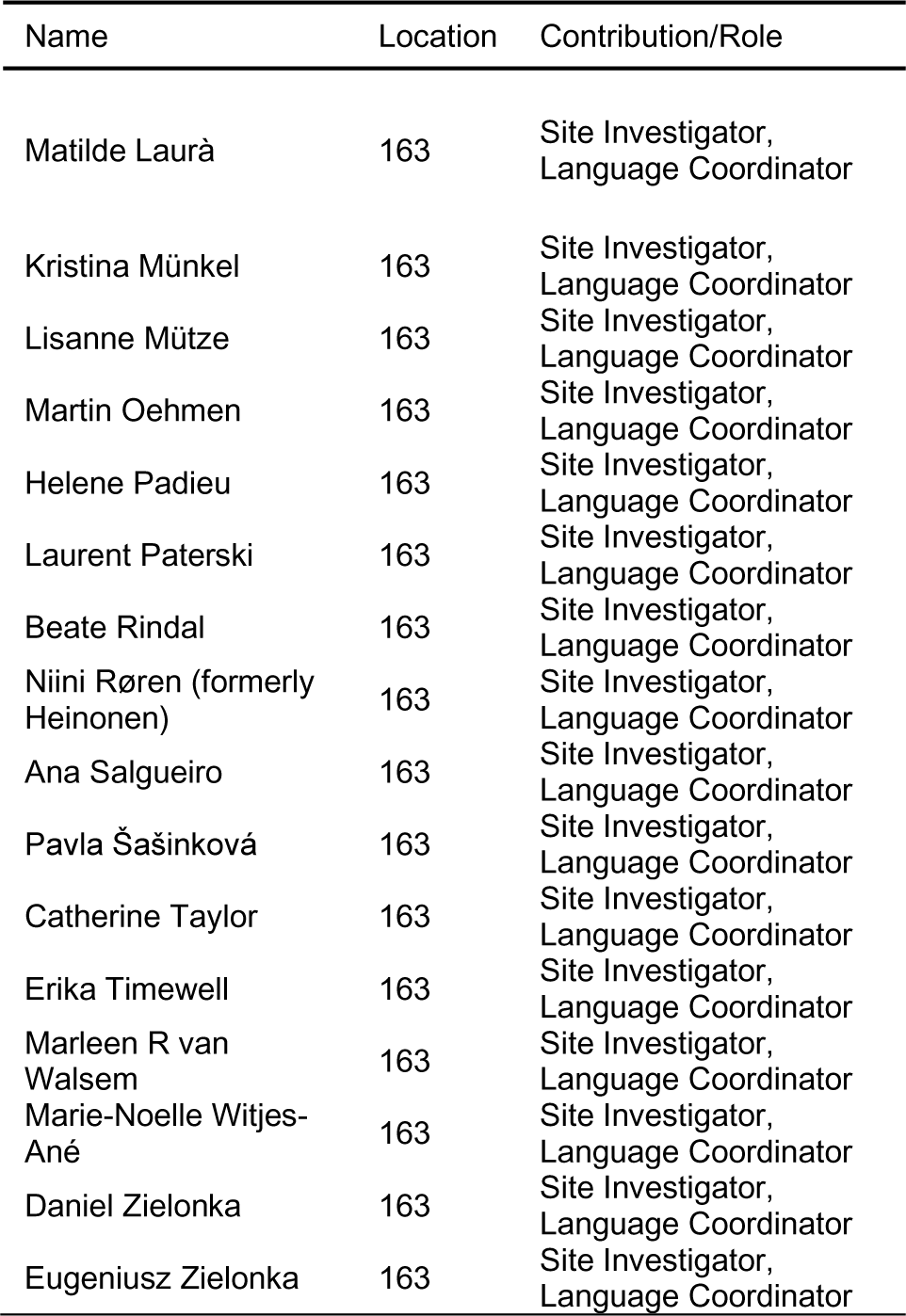

**Affiliations:**

1: Graz (Medizinische Universitäts Graz, Psychiatrie), Austria

2: Innsbruck (Universitätsklinik Innsbruck, Neurologie), Austria

3: Salzburg (Christian-Doppler-Klinik Salzburg, Universitätsklinikum der PMU, Universitätsklinik für Neurologie), Austria

4: Vienna-UNI, Austria

5: Bierbeek, Belgium

6: Bruxelles (Vrije Universiteit Brussel), Belgium

7: Bruxelles (Erasmes), Belgium

8: Bruxelles (St-Luc), Belgium

9: Charleroi (Institut de Pathologie et de Génétique (IPG)), Belgium

10: Leuven (Universitair Ziekenhuis Gasthuisberg), Belgium

11: Olomouc (Neurologická klinika, Fakultní nemocnice Olomouc), Czech Republic

12: Prague (Extrapyramidové centrum, Neurologická klinika, 1. LF UK a VFN), Czech Republic

13: Aarhus (Aarhus University Hospital), Denmark

14: Copenhagen University Hospital (Rigshospitalet, Memory clinic), Denmark

15: Odense (Odense University Hospital), Denmark

16: Aland, Finland

17: Helsinki - Vaestoliitto (Department of Medical Genetics), Finland

18: Kuopio: Anu Bruun, Finland

19: Oulu (Dep. of Neurology), Finland

20: Oulu (Dep. of Medical Genetics), Finland

21: Tampere (Terveystalo Healthcare Service Centre), Finland

22: Turku-Suvituuli (Rehabilitation Centre Suvituuli), Finland

23: Angers (Centre de référence des maladies neurogénétique-CHU d’Angers), France

24: Bordeaux (Hôpital Pellegrin), France

25: Clermont-Ferrand (Hôpital Gabriel Montpied), France

26: Creteil (Hôpital Henri Mondor), France

27: Lille-Amiens (Lille (CHRU Roger Salengro)), France

28: Lille-Amiens (Amiens (CHU Sud)), France

29: Marseille (Hôpital La Timone), France

30: Paris (Hôpital de la Pitié Salpêtrière), France

31: Rouen (Hôpital Charles Nicolle), France

32: Strasbourg (Hôpital Civil), France

33: Toulouse (Hôpital Purpan), France

34: Aachen (Universitätsklinikum Aachen, Neurologische Klinik), Germany

35: Berlin (Universitätsmedizin Berlin, Klinik und Poliklinik für Neurologie), Germany

36: Bochum (Huntington-Zentrum (NRW) Bochum im St. Josef-Hospital), Germany

37: Bremen, Germany

38: Dinslaken (Reha Zentrum in Dinslaken im Gesundheitszentrums Lang), Germany

39: Dresden (Universitätsklinikum Carl Gustav Carus an der Technischen Universität Dresden, Klinik und Poliklinik für Neurologie), Germany

40: Erlangen (Universitätsklinikum Erlangen, Molekulare Neurologie und Klinik für Neurologie), Germany

41: Freiburg (Universitätsklinik Freiburg, Neurologie), Germany

42: Hamburg (Universitätsklinikum Hamburg-Eppendorf, Klinik und Poliklinik für Neurologie), Germany

43: Hannover (Neurologische Klinik mit Klinischer Neurophysiologie, Medizinische Hochschule Hannover), Germany

44: Itzehoe (Schwerpunktpraxis Huntington Neurologie und Psychiatrie), Germany

45: Marburg KPP (Klinik für Psychiatrie und Psychotherapie Marburg-Süd), Germany

46: Marburg UNI (Universitätsklinik Marburg, Sprechstunde für choreatiforme Bewegungsstörungen), Germany

47: München (Huntington-Ambulanz im Neuro-Kopfzentrum - Klinikum rechts der Isar der Neurologischen Klinik und Poliklinik der Technischen Universität München), Germany

48: Münster (Universitätsklinikum Münster, Klinik und Poliklinik für Neurologie), Germany

49: Taufkirchen (Isar-Amper-Klinikum - Klinik Taufkirchen (Vils)), Germany

50: Ulm (Universitätsklinikum Ulm, Neurologie), Germany

51: Würzburg (Universitätsklinikum Würzburg, Neurologie), Germany

52: Bari (Neurophysiopathology of Pain Unit, Basic Medical, Neuroscience and Sensory System Department, University of Bari), Italy

53: Bologna (DIBINEM - Alma Mater Studiorum - Università di Bologna, IRCCS Istituto delle Scienze Neurologiche di Bologna), Italy

54: Brescia (Division of Biology and Genetics, Department of Molecular and Translational Medicine & Division of Neurology, Department of Clinical and Experimental Sciences, University of Brescia), Italy 55: Cagliari (Movement Disorders Center, Department of Neurology, Institute of Neurology, University of Cagliari), Italy

56: Florence (Department of NEUROFARBA, University of Florence & Careggi University Hospital, IRCSS “Don Gnocchi”), Italy

57: Genoa (Department of Neuroscience, Rehabilitation, Ophthalmology, Genetics, Maternal and Child Health, University of Genova), Italy

58: Milan (SODS Genetica delle Malattie Neurodegenerative e Metaboliche & U.O. Neurologia, Fondazione IRCCS Istituto Neurologico Carlo Besta), Italy

59: Naples (Department of Neurosciences and Reproductive and Odontostomatological Sciences Federico II University of Naples), Italy

60: Pozzilli (IS) (IRCCS Neuromed), Italy

61: IRCCS Casa Sollievo della Sofferenza, San Giovanni Rotondo, Italy

62: Rome (LIRH Foundation), Italy

63: Rome (Department of Neurology, Università Cattolica del Sacro Cuore; Institute of Translational Pharmacology & Institute of Cognitive Sciences and Technologies, National Research Council of Italy), Italy

64: Rome (Azienda Ospedaliera Sant’Andrea; Department of Neuroscience, Mental Health and Sensory Organs (NESMOS), Faculty of Medicine and Psychology, Sapienza University of Rome; Institute of Translational Pharmacology & Institute of Cognitive Sciences and Technologies, National Research Council of Italy), Italy

65: Enschede (Medisch Spectrum Twente), Netherlands

66: Groningen (Polikliniek Neurologie), Netherlands

67: Leiden (Leiden University Medical Centre (LUMC)), Netherlands

68: Maastricht, Netherlands

69: Nijmegen (Universitair Medisch Centrum St. Radboud, Neurology), Netherlands

70: Bergen (Haukeland University Hospital, Dept of Medical Genetics and Olaviken Psychiatric Hospital), Norway

71: Bergen (NKS Olaviken’s HD clinic), Norway

72: Oslo University Hospital (Dept. of Medical Genetics, Dept. of Neurology, Dept.of Neurorehabilitation), Norway

73: Trondheim (St. Olavs Hospital), Norway

74: Gdansk (St. Adalbert Hospital, Gdansk, Medical University of Gdansk, Neurological and Psychiatric Nursing Dpt.), Poland

75: Katowice (Medical University of Silesia, Katowice), Poland

76: Krakow (Krakowska Akademia Neurologii), Poland

77: Poznan (Poznan University of Medical Sciences), Poland

78: Warsaw-MU (Medical University of Warsaw, Neurology), Poland

79: Warsaw-IPiN (Institute of Psychiatry and Neurology Dep. of Genetics, First Dep. of Neurology), Poland

80: Coimbra – (Hospital Universitário de Coimbra), Portugal

81: Lisbon-Central (Hospital dos Capuchos, Centro Hositalar Lisboa Central), Portugal

82: Lisbon-HSM (Hospital de Santa Maria, Clinical Pharmacology Unit, Instituto de Medicina Molecular), Portugal

83: Lisbon-HFF (Hospital Fernando da Fonseca), Portugal

84: Porto-HGSA (Hospital Santo António-Centro Hospitalar do Porto), Portugal

85: Porto-HSJ (Hospital de São João), Portugal

86: Kazan, Russian Federation

87: Moscow – (Research Center of Neurology), Russian Federation

88: Nizhny Novgorod – (Nizhny Novgorod Medical Academy, Neurology Department), Russian Federation

89: Ufa – (Bashkir State Medical University, Department of Neurology, Neurosurgery, and Medical Genetics), Russian Federation

90: Voronezh, Russian Federation

91: Alicante-Alcoy (Hospital Virgen de los Lirios), Spain

92: Badajoz (Hospital Infanta Cristina), Spain

93: Barcelona-Bellvitge (Hospital Universitari de Bellvitge), Spain

94: Barcelona-Clínic i Provincial (Hospital Clínic i Provincial), Spain

95: Barcelona-Hospital Mútua de Terrassa, Spain

96: Barcelona-Merced (Hospital Mare de Deu de La Merced), Spain

97: Barcelona-Santa Cruz y San Pablo (Hospital de la Santa Creu i Sant Pau), Spain

98: Bilbao (Hospital de Cruces), Spain

99: Burgos (Servicio de Neurología Hospital General Yagüe), Spain

100: Canarias (Hospital Insular de Gran Canaria), Spain

101: Fuenlabrada (Hospital Universitario), Spain

102: Granada (Hospital Universitario San Cecilio, Neurología), Spain

103: Madrid-BTCIEN (Fundación CIEN), Spain

104: Madrid-Clínico (Hospital Clínico Universitario San Carlos), Spain

105: Madrid RYC (Hospital Ramón y Cajal, Neurología), Spain

106: Madrid FJD (Madrid-Fundación Jiménez Díaz), Spain

107: Murcia (Hospital Universitario Virgen de la Arrixaca), Spain

108: Oviedo (Hospital Central de Asturias), Spain

109: Palma de Mallorca (Hospital Universitario Son Espases), Spain

110: Pamplona (Complejo Hospitalario de Navarra), Spain

111: Sevilla (Hospital Universitario Virgen del Rocío), Spain

112: Sevilla (Hospital Virgen Macarena), Spain

113: Sevilla (Residencia Santa Ana), Spain

114: Valencia (Hospital la Fe), Spain

115: Zaragoza (Hospital Clínico), Spain

116: Göteborg (Sahlgrenska University Hospital), Sweden

117: Lund (Dept Neurology, Skånes Universityhospital), Sweden

118: Stockholm-Ersta, Sweden

119: Stockholm Karolinska University Hospital, Sweden

120: Umeå (Umeå University Hospital), Sweden

121: Uppsala University Hospital, Sweden

122: Bern (Swiss HD Zentrum), Switzerland

123: Bern (Zentrum für Bewegungsstörungen, Neurologische Klinik und Poliklinik, Universität Bern), Switzerland

124: Lausanne, Switzerland

125: Zürich (University Hospital and University of Zurich), Switzerland

126: Aberdeen (NHS Grampian Clinical Genetics Centre & University of Aberdeen), UK

127: Barnstaple, UK

128: Birmingham (The Barberry Centre, Dept of Psychiatry), UK

129: Blanford Forum, UK

130: Bristol (North Bristol NHs Trust, Southmead hospital), UK

131: Cambridge (Cambridge Centre for Brain Repair, Forvie Site), UK

132: Cardiff (Schools of Medicine and Biosciences, Cardiff University), UK

133: Dundee (Scottish Huntington’s Association, Ninewells Hospital), UK

134: Edinburgh (SE Scotland Genetic Service, Western General Hospital), UK

135: Exeter (Department of Neurology Royal Devon and Exeter Foundation Trust Hospital), UK

136: Fife (Scottish Huntington’s Association Whyteman’s Brae Hospital), UK

137: Forth Valley (Neurology Department, Forth Valley Royal Hospital), UK

138: Glasgow (Glasgow HD Management Clinic, Southern General Hospital), UK

139: Gloucester (Department of Neurology Gloucestershire Royal Hospital), UK

140: Hull (Castle Hill Hospital), UK

141: Launceston (Millaton Court), UK

142: Leeds (Chapel Allerton Hospital, Department of Clinical Genetics), UK

143: Leicester (Leicestershire Partnership Trust, Mill Lodge), UK

144: Liverpool (Walton Centre for Neurology and Neurosurgery), UK

145: London (St. Georges-Hospital), UK

146: London (Guy’s Hospital), UK

147: London (The National Hospital for Neurology and Neurosurgery), UK

148: London (Royal Hospital for Neuro-disability), UK

149: Manchester (Genetic Medicine, University of Manchester, Manchester Academic Health Sciences Centre and Central Manchester University Hospitals NHS Foundation Trust), UK

150: Newcastle-upon-Tyne (Centre for Life, Institute of Medical Genetics), UK

151: Northampton (St Andrew’s Healthcare), UK

152: Oxford (Oxford University Hospitals NHS Trust, Dept. of Neurosciences, University of Oxford), UK

153: Plymouth (Plymouth Huntington Disease Service, Mount Gould Hospital), UK

154: Poole (Brain Injury Service, Poole Hospital), UK

155: Preston (Neurology Department, Preston Royal Hospital), UK

156: Reading (Royal Berkshire Hospital), UK

157: Sheffield (The Royal Hallamshire Hospital– Sheffield Children’s Hospital), UK

158: Southampton (Southampton General Hospital), UK

159: Stoke on Trent (Bucknall Hospital), UK

160: Swindon (Victoria Centre, Great Western Hospital), UK

161: EHDN’s associate site in Singapore: National Neuroscience Institute Singapore,

162: Institute for Genetic and Biomedical Research, University of Milan, Italy

163: European Huntington’s Disease Network (EHDN), Ulm, Germany,

164: CHDI Foundation, Inc., New York, USA

165: 2mt Software GmbH, Ulm, Germany

166: Clinic of Neurology, Charles University and General Teaching Hospital, Prague, Czech Republic, Czech Republic

167: Center for Rare Disorders, Oslo University Hospital HF, Rikshospitalet, Norway

168: Department of Neurology, Turku University Hospital, Turku, Finland

169: Clinic of Psychiatry, Charles University and General Teaching Hospital, Prague, Czech Republic

## Appendix 3 Huntington’s disease clinical characteristics questionnaire (HD-CCQ) questions

Appendix 3 – Huntington’s disease clinical characteristics questionnaire (HD-CCQ) questions
| Symptom | Question |
| --- | --- |
| Motor | Have motor symptoms compatible with HD ever been a part of the subject’s medical history? At what age did the subject’s motor symptoms begin? |
| Cognitive | Has significant cognitive impairment (severe enough to impact on work or activities of daily living) or dementia ever been a part of the subject’s medical history? At what age did cognitive impairment first start to have an impact on daily life? |
| Apathy | Has apathy ever been a part of the subject’s medical history? At what age did apathy begin? |
| Depression | Has depression (includes treatment with antidepressants with or without a formally-stated diagnosis of depression) ever been a part of the subject’s medical history? At what age did the depression begin? |
| Perseverative/ obsessive behaviours (POB) | Has perseverative/obsessive behaviours ever been a part of the subject’s medical history? At what age did perseverative/obsessive behaviour begin? |
| Irritability | Has irritability ever been a part of the subject’s medical history? At what age did the irritability begin? |
| Violent or aggressive behaviour (VAB) | Has violent or aggressive behaviour ever been a part of the subject’s medical history? At what age did violent or aggressive behaviour begin? |
| Psychosis | Has psychosis (hallucinations or delusions) ever been a part of the subject’s medical history? At what age did psychosis (hallucinations or delusions) begin? |

Questionnaire taken from the European Huntington’s Disease Network’s (EHDN) data dictionary for Registry (Version 3).

---

**Table e-1.**
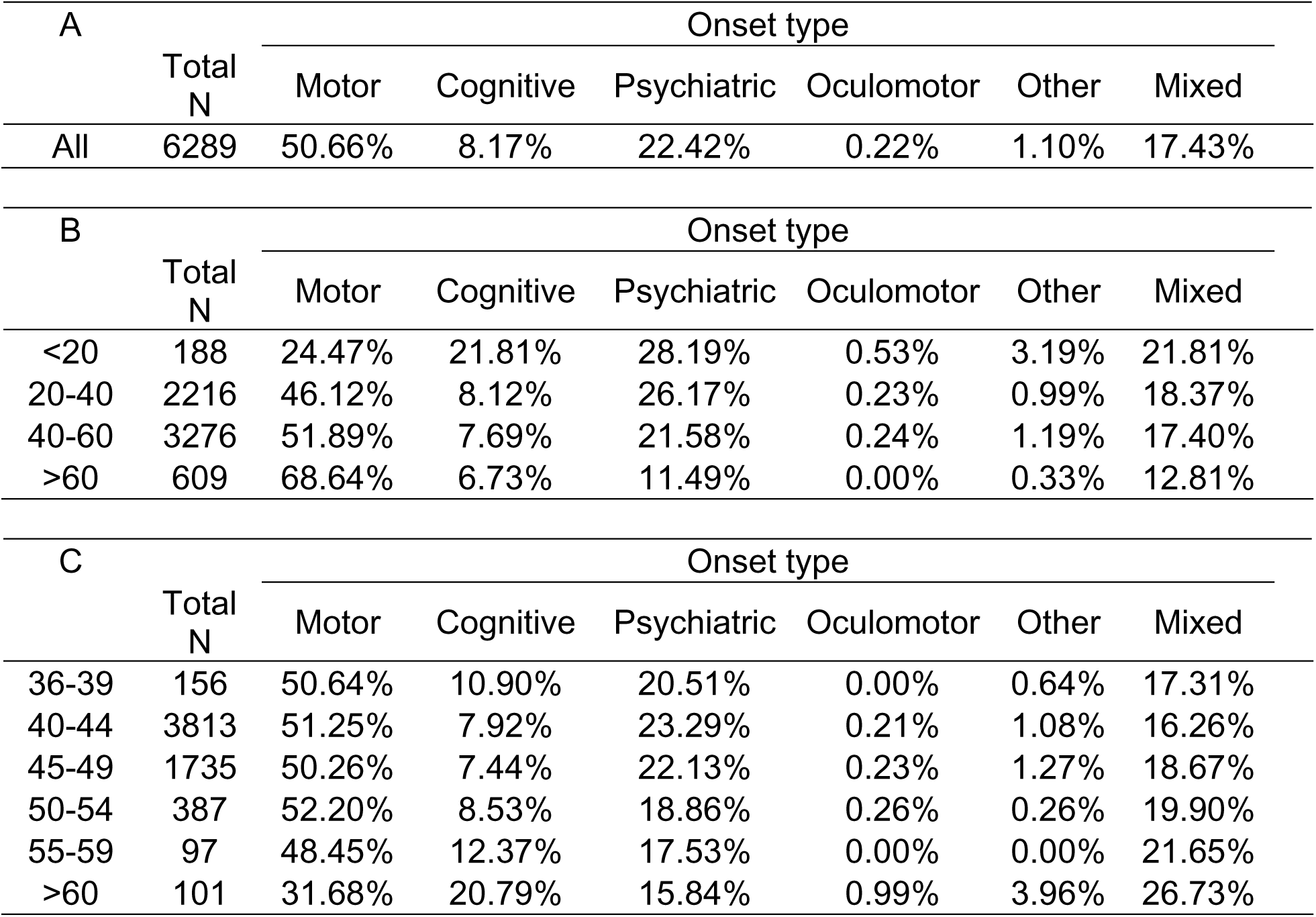
The initial manifestation of HD varies by age and CAG length. All included individuals had a pathogenic CAG length (36-93) and confirmed HD onset age determined by a rating clinician. **(A)** Percentage of different onset types in all individuals. **(B)** Percentage of different onset types observed in four age groups. **(C)** Percentage of different onset types observed across six CAG groups. These data are plotted in Fig. 1A-B.

| A | Total<br>N | Onset type |  |  |  |  |  |
| --- | --- | --- | --- | --- | --- | --- | --- |
|  |  | Motor | Cognitive | Psychiatric | Oculomotor | Other | Mixed |
| All | 6289 | 50.66% | 8.17% | 22.42% | 0.22% | 1.10% | 17.43% |

| B | Total<br>N | Onset type |  |  |  |  |  |
| --- | --- | --- | --- | --- | --- | --- | --- |
|  |  | Motor | Cognitive | Psychiatric | Oculomotor | Other | Mixed |
| <20 | 188 | 24.47% | 21.81% | 28.19% | 0.53% | 3.19% | 21.81% |
| 20-40 | 2216 | 46.12% | 8.12% | 26.17% | 0.23% | 0.99% | 18.37% |
| 40-60 | 3276 | 51.89% | 7.69% | 21.58% | 0.24% | 1.19% | 17.40% |
| >60 | 609 | 68.64% | 6.73% | 11.49% | 0.00% | 0.33% | 12.81% |

| C | Total<br>N | Onset type |  |  |  |  |  |
| --- | --- | --- | --- | --- | --- | --- | --- |
|  |  | Motor | Cognitive | Psychiatric | Oculomotor | Other | Mixed |
| 36-39 | 156 | 50.64% | 10.90% | 20.51% | 0.00% | 0.64% | 17.31% |
| 40-44 | 3813 | 51.25% | 7.92% | 23.29% | 0.21% | 1.08% | 16.26% |
| 45-49 | 1735 | 50.26% | 7.44% | 22.13% | 0.23% | 1.27% | 18.67% |
| 50-54 | 387 | 52.20% | 8.53% | 18.86% | 0.26% | 0.26% | 19.90% |
| 55-59 | 97 | 48.45% | 12.37% | 17.53% | 0.00% | 0.00% | 21.65% |
| >60 | 101 | 31.68% | 20.79% | 15.84% | 0.99% | 3.96% | 26.73% |

**Table e-2.** Age at onset, but not CAG length, is associated with initial manifestation type in adult-onset HD. **(A)** Logistic regression of initial manifestation as recorded by the clinician, coded as binary variables (1: initial manifestation of that type, 0: initial manifestation of any other type) on age at onset. **(B)** Logistic regression of initial manifestation (defined as a binary variable, as above) on CAG length. Significant associations after Bonferroni correction for 12 models are shown in bold (*P* < 4.17 x 10^-3^) and nominally significant associations in italics (*P* < 0.05). The odds ratio (OR) indicates the effect on the outcome probability associated with an increase of ten years in age. Only individuals with a confirmed onset ≥20 years, CAG 36-59 and no schizophrenia co-morbidity (ICD-10 F20, F21 or F25) were included. N=6051 for both sets of models. CI: confidence interval.

| A | Motor |  | Cognitive |  | Psychiatric |  |
| --- | --- | --- | --- | --- | --- | --- |
|  | OR (95% CI) | *P* | OR (95% CI) | *P* | OR (95% CI) | *P* |
| Age at onset | 1.06 (1.04-1.07) | **7.44 x 10<sup>-22</sup>** | 0.99 (0.99-1.00) | *3.59 x 10<sup>-2</sup>* | 0.96 (0.95-0.97) | **9.41 x 10<sup>-16</sup>** |
|  | Oculomotor |  | Other |  | Mixed |  |
|  | OR (95% CI) | *P* | OR (95% CI) | *P* | OR (95% CI) | *P* |
| Age at onset | 1.00 (1.00-1.00) | *6.06 x 10<sup>-1</sup>* | 1.00 (1.00-1.00) | *6.55 x 10<sup>-1</sup>* | 0.99 (0.98-1.00) | *2.94 x 10<sup>-2</sup>* |
| B | Motor |  | Cognitive |  | Psychiatric |  |
|  | OR (95% CI) | *P* | OR (95% CI) | *P* | OR (95% CI) | *P* |
| CAG | 1.00 (0.96-1.04) | *9.67 x 10<sup>-1</sup>* | 1.00 (0.98-1.02) | *8.05 x 10<sup>-1</sup>* | 0.96 (0.93-0.99) | *1.97 x 10<sup>-2</sup>* |
|  | Oculomotor |  | Other |  | Mixed |  |
|  | OR (95% CI) | *P* | OR (95% CI) | *P* | OR (95% CI) | *P* |
| CAG | 1.00 (1.00-1.00) | *7.93 x 10<sup>-1</sup>* | 1.00 (0.99-1.01) | *6.07 x 10<sup>-1</sup>* | 1.01 (0.99-1.04) | *9.98 x 10<sup>-3</sup>* |

**Table e-3.** The CAG repeat length of the pathogenic *HTT* allele, but not the wild-type HTT allele, is associated with the presence of motor and psychiatric symptoms in HD. Logistic regression of binary symptom data (lifetime prevalence; 0 = no symptom, 1 = reported symptom) on expanded (36-93 CAGs) and wild-type (6-35 CAGs) repeat lengths, with an interaction term. Significant associations are in bold (*P* < 2.08 x 10^-3^, correcting for number of covariates and phenotypes), and nominal associations italicised (*P* < 0.05). The odds ratio (OR) indicates the effect on the outcome probability associated with an increase of one unit in the covariate. In addition to having a confirmed onset and pathogenic CAG length, to be included individuals must have a known sex and no co-morbid diagnosis of schizophrenia, schizotypy or schizoaffective disorder. CI: confidence interval; POB: perseverative/obsessive behaviour; VAB: violent or aggressive behaviour.

|  | Motor (N=5363) |  | Cognitive (N=3141) |  | Apathy (N=2798) |  |
| --- | --- | --- | --- | --- | --- | --- |
|  | OR (95% CI) | *P* | OR (95% CI) | *P* | OR (95% CI) | *P* |
| Exp CAG | 0.94<br>(0.94 - 0.95) | **1.17 x 10<sup>-89</sup>** | 0.94<br>(0.93 - 0.95) | **7.38 x 10<sup>-51</sup>** | 0.93<br>(0.93 - 0.94) | **1.70 x 10<sup>-40</sup>** |
| WT CAG | 0.99<br>(0.98 - 1.01) | 4.55 x 10<sup>-1</sup> | 0.99<br>(0.97 - 1.01) | 2.76 x 10<sup>-1</sup> | 0.97<br>(0.95 - 0.99) | 6.98 x 10<sup>-3</sup> |
| Interaction | 1.00<br>(1.00 - 1.00) | 4.89 x 10<sup>-1</sup> | 1.00<br>(1.00 - 1.00) | 3.48 x 10<sup>-1</sup> | 1.001<br>(1.00 - 1.00) | 6.51 x 10<sup>-3</sup> |

|  | Depression (N=3439) |  | POB (N=1927) |  | Irritability (N=3165) |  |
| --- | --- | --- | --- | --- | --- | --- |
|  | OR (95% CI) | *P* | OR (95% CI) | *P* | OR (95% CI) | *P* |
| Exp CAG | 0.94<br>(0.93 - 0.95) | **1.49 x 10<sup>-19</sup>** | 0.95<br>(0.93 - 0.96) | **3.78 x 10<sup>-13</sup>** | 0.95<br>(0.94 - 0.96) | **7.37 x 10<sup>-17</sup>** |
| WT CAG | 0.99<br>(0.96 - 1.02) | 3.69 x 10<sup>-1</sup> | 0.99<br>(0.96 - 1.02) | 5.32 x 10<sup>-1</sup> | 1.02<br>(0.99 - 1.05) | 1.63 x 10<sup>-1</sup> |
| Interaction | 1.000<br>(1.00 - 1.00) | 3.26 x 10<sup>-1</sup> | 1.00<br>(1.00 - 1.00) | 6.10 x 10<sup>-1</sup> | 1.00<br>(1.00 - 1.00) | 1.67 x 10<sup>-1</sup> |

|  | VAB (N=1630) |  | Psychosis (N=597) |  |
| --- | --- | --- | --- | --- |
|  | OR (95% CI) | *P* | OR (95% CI) | *P* |
| Exp CAG | 0.94<br>(0.92 - 0.95) | **4.24 x 10<sup>-15</sup>** | 0.92<br>(0.90 - 0.95) | **5.28 x 10<sup>-8</sup>** |
| WT CAG | 0.98<br>(0.95 - 1.02) | 4.12 x 10<sup>-1</sup> | 0.94<br>(0.88 - 1.00) | 5.01 x 10<sup>-2</sup> |
| Interaction | 1.00<br>(1.00 - 1.00) | 3.67 x 10<sup>-1</sup> | 1.00<br>(1.00 - 1.00) | 4.14 x 10<sup>-2</sup> |

|  | Depressed mood<br>(N=1721) |  | Suicidal ideation<br>(N=1708) |  | Anxiety<br>(N=1717) |  | Irritability<br>(N=1727) |  |
| --- | --- | --- | --- | --- | --- | --- | --- | --- |
|  | Effect<br>(95% CI) | *P* | Effect<br>(95% CI) | *P* | Effect<br>(95% CI) | *P* | Effect<br>(95% CI) | *P* |
| *Motor* | 1.20<br>(±1.89) | 2.14X 10<sup>-1</sup> | 0.60<br>(±0.87) | 1.79X 10<sup>-1</sup> | -0.66<br>(±1.83) | 4.80X 10<sup>-1</sup> | -0.05<br>(±1.94) | 9.61X 10<sup>-1</sup> |
| *Cognitive* | -0.18<br>(±0.37) | 3.28X 10<sup>-1</sup> | -0.09<br>(±0.17) | 3.18X 10<sup>-1</sup> | -0.02<br>(±0.36) | 8.94X 10<sup>-1</sup> | 0.19<br>(±0.38) | 3.27X 10<sup>-1</sup> |
| *Apathy* | 0.48<br>(±0.37) | 1.15X 10<sup>-2</sup> | 0.12<br>(±0.17) | 1.69X 10<sup>-1</sup> | 0.62<br>(±0.36) | 8.44X 10<sup>-4</sup> | 0.39<br>(±0.38) | 4.84X 10<sup>-2</sup> |
| *Depression* | 1.70<br>(±0.37) | **2.19X 10<sup>-19</sup>** | 0.39<br>(±0.17) | **7.22x 10<sup>-6</sup>** | 0.92<br>(±0.36) | **4.54x 10<sup>-7</sup>** | -0.04<br>(±0.38) | 8.45x 10<sup>-1</sup> |
| *POB* | 0.00<br>(±0.37) | 9.97X 10<sup>-1</sup> | 0.09<br>(±0.17) | 2.95x 10<sup>-1</sup> | 0.23<br>(±0.36) | 2.13x 10<sup>-1</sup> | 0.37<br>(±0.38) | 5.33x 10<sup>-2</sup> |
| *Irritability* | 0.68<br>(±0.39) | 6.69X 10<sup>-4</sup> | 0.27<br>(±0.18) | 3.44x 10<sup>-3</sup> | 0.62<br>(±0.38) | 1.33x 10<sup>-3</sup> | 1.80<br>(±0.40) | **4.39x 10<sup>-18</sup>** |
| *VAB* | 0.07<br>(±0.41) | 7.28X 10<sup>-1</sup> | 0.00<br>(±0.19) | 9.60x 10<sup>-1</sup> | 0.22<br>(±0.40) | 2.85x 10<sup>-1</sup> | 1.01<br>(±0.42) | **2.31x 10<sup>-6</sup>** |
| *Psychosis* | -0.34<br>(±0.53) | 2.06X 10<sup>-1</sup> | 0.10<br>(±0.25) | 4.17x 10<sup>-1</sup> | -0.12<br>(±0.52) | 6.58x 10<sup>-1</sup> | -0.25<br>(±0.54) | 3.60x 10<sup>-1</sup> |
| *Age* | -0.04<br>(±0.02) | **1.98X 10<sup>-4</sup>** | -0.01<br>(±0.01) | 4.23x 10<sup>-2</sup> | -0.02<br>(±0.02) | 2.73x 10<sup>-2</sup> | -0.02<br>(±0.02) | 2.00x 10<sup>-2</sup> |
| *CAG* | -0.10<br>(±0.05) | **2.39X 10<sup>-4</sup>** | -0.03<br>(±0.02) | 1.82x 10<sup>-2</sup> | -0.07<br>(±0.05) | 5.68x 10<sup>-3</sup> | -0.01<br>(±0.05) | 7.41x 10<sup>-1</sup> |
| *Sex (F)* | 0.28<br>(±0.33) | 9.10X 10<sup>-2</sup> | -0.18<br>(±0.15) | 2.18x 10<sup>-2</sup> | 0.14<br>(±0.32) | 3.94x 10<sup>-1</sup> | 0.13<br>(±0.34) | 4.60x 10<sup>-1</sup> |
| *Duration* | -0.01<br>(±0.03) | 5.44X 10<sup>-1</sup> | 0.00<br>(±0.01) | 9.63x 10<sup>-1</sup> | -0.01<br>(±0.03) | 4.93x 10<sup>-1</sup> | 0.02<br>(±0.03) | 1.53x 10<sup>-1</sup> |

|  | Angry or aggressive<br>behaviour (N=1726) |  | Apathy (N=1719) |  | Perseverative thinking/<br>behaviours (N=1722) |  | Obsessive compulsive<br>behaviours (N=1711) |  |
| --- | --- | --- | --- | --- | --- | --- | --- | --- |
|  | Effect<br>(95% CI) | *P* | Effect<br>(95% CI) | *P* | Effect<br>(95% CI) | *P* | Effect<br>(95% CI) | *P* |
| *Motor* | 0.42<br>(±1.57) | 6.05X 10<sup>-1</sup> | 1.03<br>(±2.48) | 4.14X 10<sup>-1</sup> | 1.40<br>(±2.16) | 2.05x 10<sup>-1</sup> | 0.69<br>(±1.71) | 4.25x 10<sup>-1</sup> |
| *Cognitive* | 0.35<br>(±0.31) | 2.48X 10<sup>-2</sup> | 0.84<br>(±0.48) | 6.25X 10<sup>-4</sup> | 0.60<br>(±0.42) | 5.23x 10<sup>-3</sup> | -0.02<br>(±0.33) | 9.10x 10<sup>-1</sup> |
| *Apathy* | 0.22<br>(±0.31) | 1.63X 10<sup>-1</sup> | 3.30<br>(±0.49) | **5.15X 10<sup>-38</sup>** | 0.43<br>(±0.43) | 4.93x 10<sup>-2</sup> | 0.06<br>(±0.34) | 7.08x 10<sup>-1</sup> |
| *Depression* | 0.00<br>(±0.31) | 9.88x 10<sup>-1</sup> | 0.58<br>(±0.48) | 1.89x 10<sup>-2</sup> | 0.54<br>(±0.42) | 1.12x 10<sup>-2</sup> | 0.19<br>(±0.33) | 2.64x 10<sup>-1</sup> |
| *POB* | 0.44<br>(±0.31) | 5.29x 10<sup>-3</sup> | 0.61<br>(±0.48) | 1.35x 10<sup>-2</sup> | 2.48<br>(±0.42) | **5.56x 10<sup>-30</sup>** | 1.27<br>(±0.33) | **9.69x 10<sup>-14</sup>** |
| *Irritability* | 0.52<br>(±0.33) | 1.88x 10<sup>-3</sup> | -0.01<br>(±0.51) | 9.80x 10<sup>-1</sup> | -0.06<br>(±0.45) | 7.82x 10<sup>-1</sup> | 0.04<br>(±0.35) | 8.13x 10<sup>-1</sup> |
| *VAB* | 1.51<br>(±0.34) | **8.81x 10<sup>-18</sup>** | 0.76<br>(±0.54) | 5.58x 10<sup>-3</sup> | 0.25<br>(±0.47) | 2.99x 10<sup>-1</sup> | 0.13<br>(±0.37) | 4.85x 10<sup>-1</sup> |
| *Psychosis* | 0.21<br>(±0.44) | 3.63x 10<sup>-1</sup> | 1.23<br>(±0.70) | 5.50x 10<sup>-4</sup> | 0.94<br>(±0.61) | 2.49x 10<sup>-3</sup> | 0.13<br>(±0.48) | 6.10x 10<sup>-1</sup> |
| *Age* | -0.01<br>(±0.02) | 2.21x 10<sup>-1</sup> | 0.03<br>(±0.03) | 6.18x 10<sup>-2</sup> | 0.01<br>(±0.02) | 2.95x 10<sup>-1</sup> | -0.04<br>(±0.02) | **1.54x 10<sup>-4</sup>** |
| *CAG* | -0.01<br>(±0.04) | 5.57x 10<sup>-1</sup> | 0.05<br>(±0.07) | 1.26x 10<sup>-1</sup> | 0.07<br>(±0.06) | 3.61x 10<sup>-2</sup> | -0.07<br>(±0.05) | 6.39x 10<sup>-3</sup> |
| *Sex (F)* | 0.01<br>(±0.27) | 9.47x 10<sup>-1</sup> | -0.47<br>(±0.43) | 3.45x 10<sup>-2</sup> | -0.12<br>(±0.38) | 5.34x 10<sup>-1</sup> | -0.26<br>(±0.30) | 8.45x 10<sup>-2</sup> |
| *Duration* | 0.02<br>(±0.03) | 1.75x 10<sup>-1</sup> | 0.05<br>(±0.04) | 1.84x 10<sup>-2</sup> | 0.04<br>(±0.04) | 4.62x 10<sup>-2</sup> | 0.04<br>(±0.03) | 8.24x 10<sup>-3</sup> |

**Table e-4.** Scores for neuropsychiatric symptoms from the short-form Problem Behaviors Assessment questionnaire (PBA-s) correlate with specific responses from the HD clinical characteristics questionnaire (HD-CCQ). Generalised linear models regressed the product of the frequency and severity scores from PBA-s questions against binary lifetime prevalence data on 8 symptoms from the HD Clinical Characteristics Questionnaire (italics, column 1) and other covariates. For binary covariates (CCQ symptoms and sex) “effect” is the increase/decrease in the clinical score associated with presence of that covariate. For quantitative covariates (age, CAG, duration), “effect” is the change in clinical score associated with an increase of one unit in the covariate. In addition to having a confirmed onset and pathogenic CAG length (36-93), individuals must have no co-morbid diagnosis of schizophrenia, schizotypy or schizoaffective disorder. Significant associations after Bonferroni correction for 11 phenotypes and 12 covariates are shown in bold (*P* < 3.79 x 10^-4^) and nominally significant *P* values are italicised (*P* < 0.05). CI: confidence interval; POB: perseverative/ obsessive behaviour; VAB: violent or aggressive behaviour; Sex (F): female.

|  | Delusions/paranoid thinking (N=1708) |  | Hallucinations (N=1710) |  | Disorientated behaviour (N=1701) |  |
| --- | --- | --- | --- | --- | --- | --- |
|  | Effect (95% CI) | *P* | Effect (95% CI) | *P* | Effect (95% CI) | *P* |
| *Motor* | 0.20<br>(±0.95) | 6.87x 10<sup>-1</sup> | 0.02<br>(±0.54) | 9.31x 10<sup>-1</sup> | 0.67<br>(±1.48) | 3.73x 10<sup>-1</sup> |
| *Cognitive* | -0.03<br>(±0.18) | 7.16x 10<sup>-1</sup> | 0.01<br>(±0.10) | 8.22x 10<sup>-1</sup> | 0.80<br>(±0.29) | **6.53x 10<sup>-8</sup>** |
| *Apathy* | 0.00<br>(±0.18) | 9.85x 10<sup>-1</sup> | 0.09<br>(±0.11) | 9.44x 10<sup>-2</sup> | 0.20<br>(±0.29) | 1.93x 10<sup>-1</sup> |
| *Depression* | 0.16<br>(±0.18) | 7.10x 10<sup>-2</sup> | 0.05<br>(±0.10) | 3.97x 10<sup>-1</sup> | 0.08<br>(±0.29) | 5.73x 10<sup>-1</sup> |
| *POB* | 0.03<br>(±0.18) | 7.35x 10<sup>-1</sup> | -0.04<br>(±0.10) | 4.58x 10<sup>-1</sup> | 0.45<br>(±0.29) | 2.13x 10<sup>-3</sup> |
| *Irritability* | 0.11<br>(±0.19) | 2.42x 10<sup>-1</sup> | -0.10<br>(±0.11) | 6.80x 10<sup>-2</sup> | -0.21<br>(±0.31) | 1.79x 10<sup>-1</sup> |
| *VAB* | 0.00<br>(±0.20) | 9.94x 10<sup>-1</sup> | -0.01<br>(±0.12) | 9.16x 10<sup>-1</sup> | 0.59<br>(±0.32) | **3.44x 10<sup>-4</sup>** |
| *Psychosis* | 1.19<br>(±0.26) | **5.56x 10<sup>-19</sup>** | 0.81<br>(±0.15) | **4.67x 10<sup>-25</sup>** | 0.68<br>(±0.42) | 1.60x 10<sup>-3</sup> |
| *Age* | -0.01<br>(±0.01) | 3.08x 10<sup>-1</sup> | 0.00<br>(±0.01) | 5.61x 10<sup>-1</sup> | 0.03<br>(±0.02) | 8.23x 10<sup>-4</sup> |
| *CAG* | -0.01<br>(±0.03) | 2.81x 10<sup>-1</sup> | 0.00<br>(±0.02) | 6.33x 10<sup>-1</sup> | 0.03<br>(±0.04) | 1.28x 10<sup>-1</sup> |
| *Sex (F)* | 0.07<br>(±0.16) | 4.19x 10<sup>-1</sup> | -0.08<br>(±0.09) | 9.84x 10<sup>-2</sup> | -0.07<br>(±0.26) | 5.75x 10<sup>-1</sup> |
| *Duration* | 0.02<br>(±0.02) | 2.81x 10<sup>-2</sup> | 0.00<br>(±0.01) | 2.92x 10<sup>-1</sup> | 0.05<br>(±0.03) | **2.92x 10<sup>-4</sup>** |

**Figure e-1.**
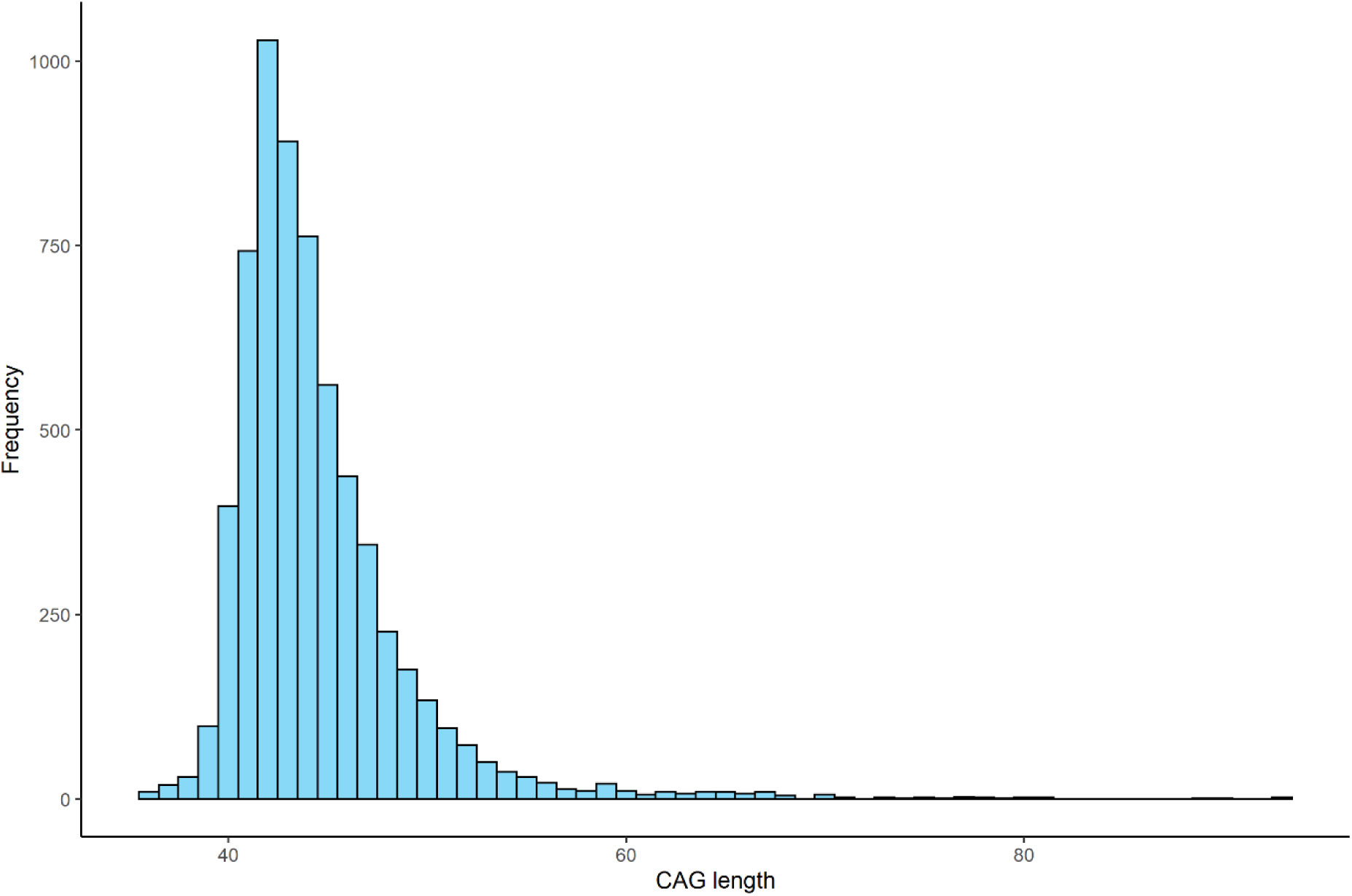
Histogram of pure CAG repeat lengths against frequency for the Registry population analysed in this study. Shown are *HTT* CAG length frequencies for the largest pathogenic CAG (36-93) in individuals with clinical onset of HD (N=6316).

**Figure e-2.**
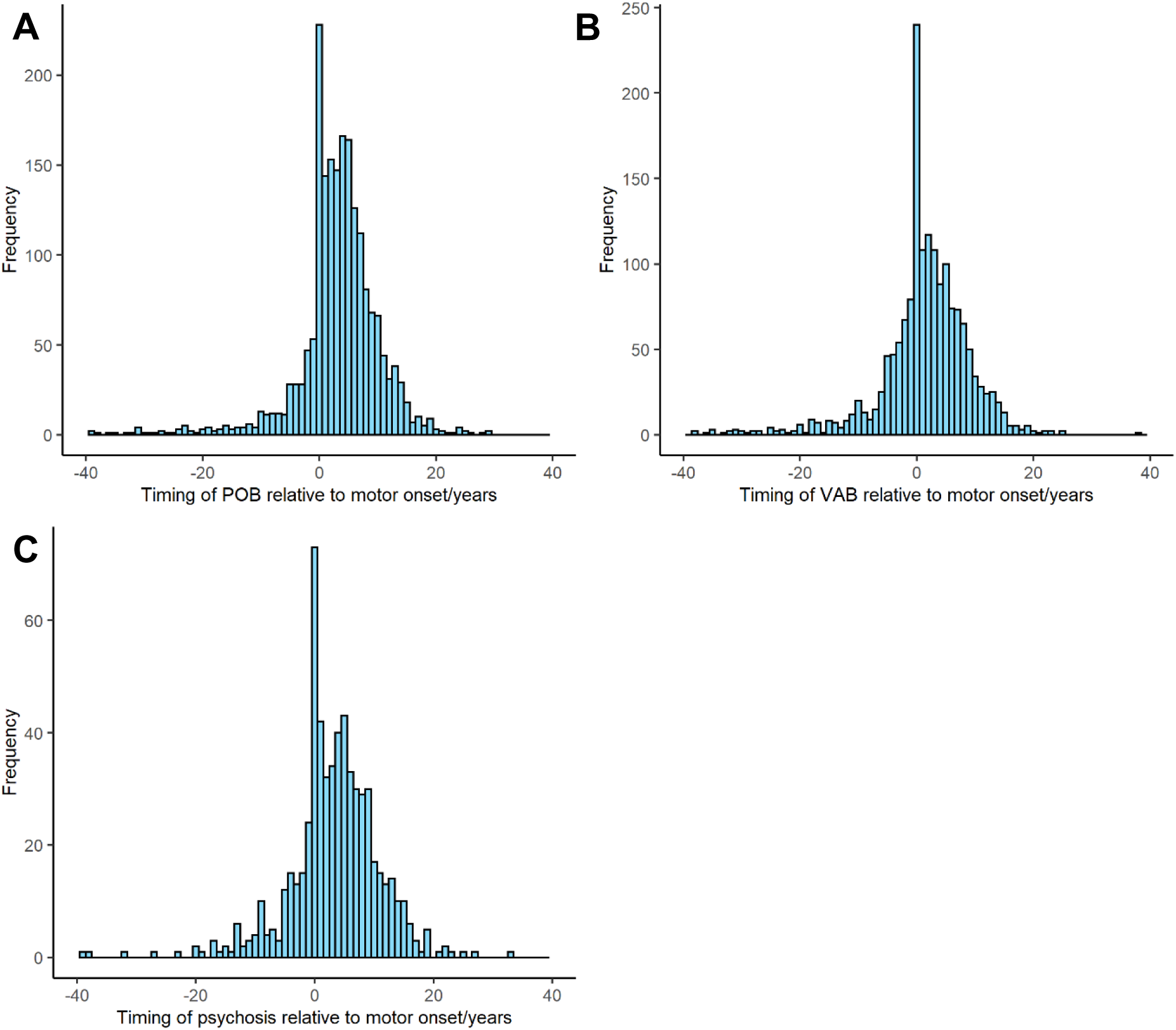
The onsets of cognitive and psychiatric symptoms relative to motor onset in HD. The age at onset of motor symptoms was subtracted from the age at onset of each cognitive/psychiatric symptom when present. Timings of up to +/− 40 years relative to motor onset shown. Only individuals with a rater-confirmed age at onset and CAG length (36-93) were included. Data from HD-CCQ. (**A**) Perseverative/obsessive behaviour (POB) N=1973, skew = −1.54; (**B**) Violent or aggressive behaviour (VAB) N=1663, skew = −1.30; (**C**) Psychosis N=619, skew = - 0.46. Only individuals with a known age at onset and CAG length (36-93) were included.

